# Bayesian Pharmacometrics Analysis of Baclofen for Alcohol Use Disorder

**DOI:** 10.1101/2022.10.25.513675

**Authors:** Nina Baldy, Meysam Hashemi, Nicolas Simon, Viktor K. Jirsa

## Abstract

Alcohol use disorder (AUD) also called alcohol dependence is a major public health problem, which affects almost 10% of the world’s population. Baclofen as a selective GABA_B_ receptor agonist has emerged as a promising drug for the treatment of AUD, however, its optimal dosage varies according to individuals, and its exposure-response relationship has not been well established yet. In this study, we use a principled Bayesian workflow to estimate the parameters of a pharmacokinetic (PK) population model from Baclofen administration to patients with AUD. By monitoring various convergence diagnostics, the probabilistic methodology is first validated on synthetic longitudinal datasets and then, applied to infer the PK model parameters based on the clinical data that were retrospectively collected from outpatients treated with oral Baclofen. We show that state-of-the-art advances in automatic Bayesian inference using self-tuning Hamiltonian Monte Carlo (HMC) algorithms with a leveraged level of information in priors provide accurate predictions on Baclofen plasma concentration in individuals. This approach may pave the way to render non-parametric HMC sampling methods sufficiently easy and reliable to use in clinical schedules for personalized treatment of AUD.

## 1. Introduction

Alcohol use disorder (AUD) is a major public health problem, which affects almost 10% of the world’s population (Schuckit, 2009). The harmful use of alcohol is responsible for 5.1% of the global burden of disease, results in more than 200 diseases, injuries and other health conditions, and eventually contributes to 3 million deaths globally each year (i.e., 5.3% of all deaths according to World Health Organization (2022)). Beyond health consequences, this brain disorder brings significant mental and behavioral symptoms, as well as social and economic losses to individuals and society at a large scale.

Baclofen is a selective agonist of the gamma-aminobutyric acid B receptors (GABA_B_), which may exert an inhibitory action on dopaminergic neurons (Doherty and Gratton, 2007). It was originally marketed as a muscle relaxant for the treatment of neurological-induced spasticity. Various experiments have shown a positive effect of Baclofen on AUD in rodents and non-human primates (see Colombo and Gessa (2018) for a review). Although there is no clear evidence on the dosing, efficacy, safety, and ideal duration of Baclofen treatment for AUD, clinical studies have shown its promising medication in patients with moderate to severe AUD (Addolorato et al., 2002; Garbutt et al., 2021). Preclinical experiments have also shown its efficacy in reducing alcohol withdrawal syndrome (Brennan et al., 2013) and voluntary alcohol intake (Colombo et al., 2004). The optimal dosage of Baclofen varies according to individuals and has not been well established with a general agreement. It is mainly prescribed to patients for whom the existing treatments have not been effective, or in those with liver disease due to its minimal damage to the liver (see de Beaurepaire et al. (2019) for a review of the use of Baclofen as a treatment in human AUD). Although in recent years, Baclofen has been used to reduce craving, voluntary alcohol intake and withdrawal syndrome of alcoholic patients, a wide inter-individual variability has been observed, and the potential high risk of sedation is unknown (Imbert et al., 2015; Simon et al., 2018). Thus, the precise dosing efficacy and the ideal duration of treatment in different AUD patient groups need to be further evaluated.

Mathematical modeling is widely used in many areas of science to learn about the data generation process, make predictions on outcomes (unseen data), and to justify hypotheses. In the modeling framework, differential equations provide us with the natural evolution of the system under study at any given time point. They are often used to describe the principles that govern the dynamics of the system that allow us to adequately describe the processes involved, and make quantitative predictions, for instance on the rate of change of drug concentration and clinical efficacy.

In this study, we use a pharmacokinetic (PK) population model of the Baclofen effect on patients with AUD to find the causal relationship between dose and exposure. The PK analysis has been widely used in clinical development with several applications, for instance, to improve the understanding of in vivo behavior of the complex delivery systems as it allows for the separation of the drug-, carrier-, and pharmacological system-specific parameters (Meibohm and Derendorf, 1997; Upton and Mould, 2014; Sheiner and Ludden, 1992; Bonate, 2011). PK analysis is currently an indispensable component of drug discovery by ensuring that robust evidence from preclinical models closely shapes the design of clinical studies (Zou et al., 2020). Although PK modeling has facilitated the drug development process, the accurate and reliable estimation of its parameters from noisy data is a major challenge for conducting clinical research to determine the safety and efficacy of a drug within a particular disease or specific patient population. To estimate unknown quantities, optimization methods (within Frequentist approach) are often used in practice by defining an objective (or a cost) function to score the performance of the model by comparing the observed with the predicted values. However, such a parametric approach results in only a point estimation, and the optimization algorithms may easily get stuck in a local maximum, requiring multi-start strategies to address the potential multi-modalities. Moreover, the estimation depends critically on the form of objective function defined for optimization, and the models involving differential equations often have non-identifiable parameters (Hashemi et al., 2018).

In this study, we use the Bayesian approach to address these challenges in the estimation of the PK population model parameters from synthetic data (generated by known values for validation), and then routine clinical data that were retrospectively collected from 67 adult outpatients treated with oral Baclofen. The Bayesian framework is a principled method for inference and prediction with a broad range of applications, while the uncertainty in parameter estimation is naturally quantified through probability distribution placed on the parameters updated with the information provided by data (Gelman et al., 2014a; Bishop, 2006). Such a probabilistic technique provides the full posterior distribution of unknown quantities in the underlying data generating process given only observed responses and the existing information about uncertain quantities expressed as prior probability distribution (Ferreira et al., 2020; Hashemi et al., 2021). In other words, Bayesian inference provides all plausible parameter ranges consistent with observation by integrating the information from both domain expertise and the experimental data. In the context of clinical trials, using Frequentist approach, prior information (based on evidence from previous trials) is utilized only in the design of a trial but not in the analysis of the data (Jack Lee and Chu, 2012; Gupta, 2012). On the other hand, Bayesian approach provides a formal mathematical framework to combine prior information with available information at the design stage, during the conduct of the experiments, and at the data analysis stage (Spiegelhalter et al., 1999; Berry, 2006; Yarnell et al., 2021). To conduct Bayesian data analysis, Markov chain Monte Carlo (MCMC) methods have often been used to sample from and hence, approximate the exact posterior distributions. However, MCMC sampling in high-dimensional parameter spaces, which converge to the desired target distribution, is non-trivial and computationally expensive (Betancourt et al., 2014; Betancourt, 2017). In particular, the use of differential equations (such as PK population models) together with noise in data raise many convergence issues (Hashemi et al., 2020; Grinsztajn et al., 2021; Jha et al., 2022). Designing an efficient MCMC sampler to perform principled Bayesian inference on high-dimensional and correlated parameters remains a challenging task. Although the Bayesian inference requires painstaking model-specific derivations and hyper-parameter tuning, probabilistic programming languages such as Stan (Carpenter et al., 2017) provide high-level tools to reliably solve complex parameter estimation problems. Stan (see https://mc-stan.org) is a state-of-the-art platform for high-performance statistical computation and automatic Bayesian data analysis, which provides advanced algorithms (Hoffman and Gelman, 2014), efficient gradient computation (Margossian, 2018), and is enriched with numerous diagnostics to check whether the inference is reliable (Vehtari et al., 2021).

In the present work, to estimate the posterior distribution of PK model parameters with MCMC, we use a self-tuning variant of Hamiltonian Monte Carlo (HMC) in Stan. This algorithm adaptively tunes the HMC’s parameters, making the sampling strategy more efficient and automatic (Hoffman and Gelman, 2014). The principled Bayesian setting (Gelman et al., 2020) in this study-validated on synthetic data- enables us to efficiently and accurately estimate the Baclofen effect on patients with AUD. This work may pave the way to reliably predict the treatment drug efficacy from longitudinal patient data, optimizing strategies for clinical decision, especially in brain disorders.

## 2. Materials and methods

### 2.1. Clinical data

In this study, we used the routine clinical data retrospectively collected from 67 adult outpatients with AUD treated with oral Baclofen in the Department of Addictology at Sainte Marguerite Hospital, Marseille, France. Ethics committee approval and patient consent are not compulsory in France to use retrospective therapeutic drug monitoring data, so no informed consent had to be collected. All patients met the DSM-5 criteria for AUD (American Psychiatric Association, 2013). The data has been thoroughly described and published in a previous study by Imbert et al. (2015). The clinical event schedule (dosing and measurements) of each subject is specified according to the Nonlinear Mixed Effects Modeling software (NONMEM, (Beal et al., 2009)) conventions. The number of observations is subject-dependant, with a total of 427 recordings. For each patient, the number of measurements is comprised of between 1 and 16 observations (median = 6). The total follow-up time ranges between 1 and 613 days (average is 185 days).

### 2.2. Synthetic data

Using synthetic data for fitting allows us to validate the inference process as we know the ground-truth of the parameters being estimated. Therefore, we can use standard error metrics to measure the similarity between the inferred parameters and those used for data generation. Synthetic data were generated following the same event schedule as empirical clinical data and using a one-compartment population model with first-order absorption (see Eq. (1)). The synthetic data was generated using the R package mrgsolve (Baron, 2022), which enables simulation from ODE-based hierarchical models, including random effects and covariates. The mrgsolve package (Elmokadem et al., 2019) uses Livermore Solver for Ordinary Differential Equations (LSODE; LSO) of the Fortran ODEPACK library (Hindmarsh, 1992) integrated in R through the Rcpp (Eddelbuettel and Francois, 2011) package.

### 2.3. Population model for pharmacometrics

Pharmacokinetic (PK) models are currently used to predict clinical response in humans through understanding the mechanism of action and disease behavior for a given drug. Inference on the PK model from data can optimize the design of clinical trials, provide the appropriate dose range, anticipate the effect in certain subpopulations, and better predict drug-drug interactions. The PK population models invoke nonlinear mixed-effects models that allow quantifying exposure-response relationships on both global population and individual levels (Bonate, 2011; Owen, 2014; Keizer et al., 2018). Such a modeling approach provides a flexible tool for hypothesis testing and refining the assumption, e.g., the random phenomena underlying the exposure-response mechanism by including group-specific (in the present case, subject-specific) effects within a population. This model also allows the inclusion of covariates whose influence on the inter-group variations can be inferred.

In the present work, drug concentration in the plasma is modeled by a one-compartment model with first-order absorption

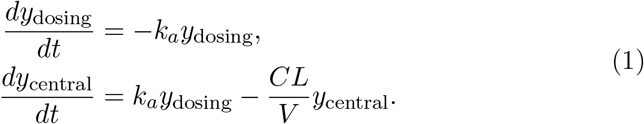

The above linear ODE system describes the evolution of the amount *y* of the drug in both dosing compartment^1^ and the central compartment (plasma). First order kinetics parameter *k*_*a*_ (h^−1^) is the absorption constant of the drug. The parameter *V* (L) is the volume of distribution, which represents the hypothetical volume in which the drug would have to be diluted at the same level as in plasma. The clearance *CL* (L/h) controls the elimination of the drug from the organism. Such a linear ODE system has an analytical solution as we used in this study. Let *y* = (*y*_dosing_, *y*_*central*_)^*T*^, the linear system given by Eq. (1) can be written in matrix form

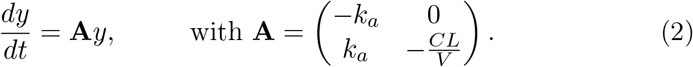

Solutions of this homogeneous linear system at time *t* are given by

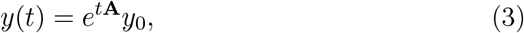

where *y*_0_ = *y*(0) is the initial condition and *e*^*t***A**^ denotes the matrix exponential^2^.

### 2.4. Bayesian modeling

In Bayesian modeling, all model parameters are treated as random variables and the values vary based on the underlying probability distribution (Bishop, 2006). That is, in the case of a one-compartment PK model, kinetic parameters *CL*, *V* and *k*_*a*_ will be interpreted as random variables, and we aim to infer their probability distributions based on *prior* knowledge (e.g., derived from physiological information or previous evidence) updated with available information in the observed data through the so-called *likelihood* function, i.e. the probability of some observed outcomes given a set of parameters. Although the likelihood function can provide the best-fit points (maximum likelihood estimators), we are interested in the whole posterior distribution over parameters as it contains all relevant information about parameters after observing data to perform inference and prediction.

Formally, given data *y* and model parameters *θ*, Bayes rule gives the posterior distribution as

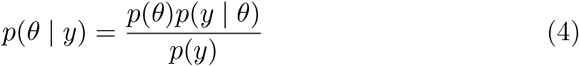

that combines and actualizes prior knowledge on parameters (before seeing data) with knowledge acquired from observed data (through likelihood function). The denominator 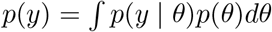 represents the probability of the data and it is known as evidence or marginal likelihood. In the context of inference, this term amounts to simply a normalization factor, thus Eq. (4) is reduced to a proportionality relation

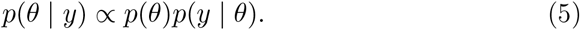

The Bayesian approach applied to PK analysis provides a fully probabilistic description of unknown quantities that allows not only a straightforward interpretation of the inferred parameters and outcomes, but also the modeling of uncertainty about the inferred values of these quantities. Moreover, it provides us with a principled method for model comparison, selection, and decision-making by measuring the out-of-sample model predictive accuracy (i.e., the measure of the model’s ability in new data prediction). To assess the predictive accuracy, we used Watanabe-Akaike (or widely available) information criterion (WAIC; Gelman et al. (2014b)). This metric is a fully Bayesian information criterion that uses the whole posterior distribution, thus enabling us to integrate our prior knowledge in the model selection. Following Gelman et al. (2014b), WAIC is given by:

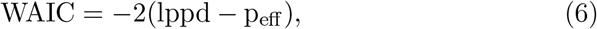

where *lppd* is the log pointwise predictive density for a new data point (as the accuracy term ^3^), and *p*_*eff*_ is the effective number of parameters (as penalty term to adjust for overfitting) ^4^. In practice, we can replace the expectations with the average over the draws from the full posterior to calculate WAIC (for more details see Gelman et al. (2014b)). Note that WAIC uses the whole posterior distribution rather than the point estimation used in the classical information criteria such as AIC and BIC. Finally, the relative difference in WAIC is used to measure the level or the strength of evidence for each candidate model under consideration. The lower value of WAIC indicates a better model fit. Following Burnham and Anderson (2002), a relative difference larger than 10 between the best model (with minimum WAIC) and other candidate models indicates that an alternative model is very unlikely (i.e., an alternative model has essentially no support).

### 2.5. MCMC sampling

Monte Carlo (MC) sampling is a family of computational algorithms for uncertainty quantification by drawing random samples from distributions, in which the sampling process does not require knowledge of the whole distribution. Markov chain Monte Carlo (MCMC) methods construct Markov chains (sequences of probabilistically linked states with the probability of transitioning to a given state depending only on the current state) that have a desired stationary probability distribution to extract samples from this distribution (Gelman et al., 2014a). Using MCMC, the samples are obtained by retrieving the history of states visited by the chain; if the chain is run long enough, these samples will originate from the stationary distribution (independent of starting states). From these samples, we can construct Monte Carlo estimates describing the target probability distribution such as mean and quantiles (i.e., the sample average approximates an expectation with respect to the stationary distribution of the chain). In this context, Monte Carlo standard errors (MCSE) provide an indication of the quality of reported estimates, i.e., a quantification of the estimation noise. MCSE of an estimator 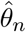 is given by the posterior standard deviation divided by the square root of the effective sample size. In addition to sample mean and sample standard deviation, MCMC methods provide `MCSE` the estimated error (standard deviation) in the posterior mean estimation, on the scale of the parameter value (Carpenter et al., 2017).

### 2.6. Adaptive HMC sampling using Stan

Stan (https://mc-stan.org) is an open-source statistical tool that implements automatic gradient-based algorithms for Bayesian modeling and probabilistic machine learning (Carpenter et al., 2017). Hamiltonian Monte Carlo (HMC) is a powerful MCMC method that uses the derivatives of the density function being sampled to generate efficient transitions exploring the whole posterior distributions in complex probabilistic models (Duane et al., 1987; Neal, 2010). However, the performance of HMC is highly sensitive to the algorithmic parameters (step size and the number of steps in the leapfrog integrator). Stan provides a self-tuning variant of the HMC, the so-called No U-Turn Sampler (Hoffman and Gelman, 2014), which uses a recursive algorithm to eliminate the need to set the hyperparameters i.e., it adaptively tunes the HMC algorithm without requiring the user intervention (Hoffman and Gelman, 2014; Betancourt, 2017). The NUTS algorithm avoids the random walk behavior and sensitivity to correlated parameters to construct efficient Markov chain exploration of the distribution’s *typical set*, that is, the set concentrating the majority of the volume-density trade-off (Betancourt, 2017).

Starting from an initial value (random or defined by the user), the NUTS algorithm updates parameters *θ* through a series of iterations by evolving them according to Hamiltonian dynamics and submitting them to a Metropolis proposal. More precisely, HMC transforms the problem of sampling from a target distribution into the problem of simulating Hamiltonian dynamics; it artificially introduces an auxiliary momentum *ρ* (to suppress such random walk behavior), which extends the representation of the target parameter space into a phase space of joint parameters (*ρ*, *θ*). The auxiliary momentum is sampled from the conditional distribution *π*(*ρ*|*θ*). In Stan implementation, it is sampled from a multivariate normal distribution 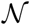 independent of *θ*,

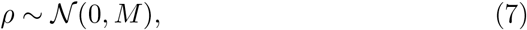

with covariance matrix *M* estimated as the inverse of posterior covariance during a *warm-up* phase. Joint phase space parameters (*ρ*, *θ*) are evolved through Hamiltonian dynamics (expression of the Hamiltonian is given in Appendix A.1) by solving the Hamiltonian equations of motion (see Appendix A.2). To do so, Stan uses the leapfrog integrator (see Appendix A.3). The output of the integrator (*ρ*^*^, *θ*^*^) is then submitted as a move proposal inside the Metropolis algorithm and is either accepted or rejected with a given probability and the system is evolved accordingly. The process is repeated iteratively starting with the momentum (re)sampling step (Eq. (7)).

### 2.7. Convergence of MCMC sampling

After running a class of MCMC algorithms, it is necessary to monitor the convergence of the samples. This can be carried out in different ways including traceplots (evolution of parameter estimates from MCMC draws over the iterations), pair plots (to identify collinearity between variables), and autocorrelation plots (to measure the degree of correlation between draws of MCMC samples). More quantitative metrics are also used in this study to assess the MCMC convergence as described in the following.

*Split-*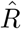 - Sampling from target distribution to estimate expectations can only be reliable if the chains have converged to the stationary distribution. The primary step to ensure the reliability of MCMC posterior chains is to check the convergence of chains with different random initializations. MCMC is asymptotically exact in the limit of infinite runs (*N* → ∞), however, in our finite-time settings, MCMC convergence cannot be guaranteed by a limited number of samples, thus, we have to rely on diagnostic quantities to assure the convergence. Potential scale reduction statistic 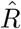 (Gelman and Rubin, 1992) is widely used in statistics to assess the convergence of the multiple Markov chains involved in posterior sampling. The 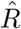 diagnostic provides an estimate of how much variance could be reduced by running the chains longer. 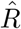 close to 1 (in practice, lower than 1.1) indicates good mixing of the chains; otherwise, the chains have not converged to the same stationary distribution. Computation of 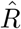 relies on between- (*B*) and within- (*W*) chain variances^5^. In particular, the marginal posterior variance is (over-) estimated as a combination of these two quantities

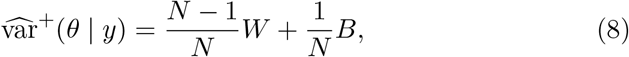

where *N* is the number of draws in each of *M* chains. The potential scale reduction 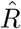 is then defined as

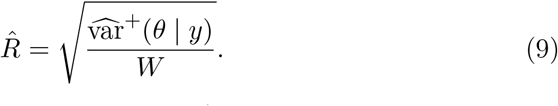

Stan reports summary statistics including 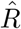 computed on the split-half chains, an additional precaution to detect non-stationarity in individual chains^6^.

#### Divergent transitions

The trajectories generated by numerical (leapfrog) integrator in the Hamiltonian equations may diverge, typically due to highly varying posterior curvature (such as funnel-shaped distributions; Betancourt (2017)), then, the samples cannot be trusted. With symplectic integrators^7^, divergences build up very quickly which makes them easy to detect. We have carefully monitored the Stan summary statistics for divergent transitions encountered during sampling.

#### Effective sample size

A complication with MCMC methods is the auto-correlation within the generated samples of a chain, often caused by strong correlations among variables. Higher autocorrelation in chains increases the uncertainty of the estimation of posterior quantities of interest. Highly correlated MCMC samplers affect the efficiency of the sampler as it requires more samples to produce the same level of Monte Carlo error for an estimate (Gelman et al., 2014a). Effective sample size (ESS) provides an estimation of the independency of draws from posterior distributions that generated Markov chain would be equivalent to. Thus, a large enough ESS is an important factor for the reliability of the inferred quantities from posterior draws. We use the bulk-ESS and tail-ESS computed on rank-normalized posterior draws (Vehtari et al., 2021) to assess the efficiency of sampling in the bulk (mean estimates efficiency), and tail (quantiles estimates efficiency) of the sampled posterior distribution. In the case of sampling with four individual chains (with *iter* = 2000 and *warmup* = 1000), we consider a minimum ESS of 400 (Vehtari et al., 2021).

#### Posterior predictive checks

Validating model assumptions and assessing the reliability of the model performance requires both domain expertise and the evaluation of the model’s predictive performance. A Posterior predictive check (PPC) is often used for model validation, i.e., generating data from the model using the parameters drawn from the estimated posterior, then comparing it with observed data. PPCs address such questions as does the fitted model and its estimated parameters generate data similar to those observed experimentally? Are the individual and population variability such as the influence of covariates consistently modeled? The principle of PPCs is to simulate multiple sets of data according to the predictive distribution *y*_*i*_ ~ *y*^*rep*^, *i* = 1, 2, · · ·, *N*, and compare them to the observed data *y*, visually or quantified by summary statistics. Visual predictive checks usually depict the predicted mean and 95% confidence/credibility intervals (2.5% and 97.5% empirical quantiles of the posterior predictive draws) confronted with observed data. Note that PPCs are not predictive per se because we are not comparing predicted data to new observations but rather we compare it to the same data that the model was fitted with. Such a process involves double use of the data and is not to be confused with validation prediction which is evaluated on a separate validation dataset or unseen data (e.g., using WAIC). Systematic discrepancies between model predictions and data would indicate an inadequate model predictive power and in this case, could be investigated to identify model failures.

### 2.8. Solving ODE systems in Stan

Linear ODE systems have the virtue of being analytically solvable, which not only provides an exact solution but is also very computationally efficient. The one-compartment linear ODE system (Eq.(1)) was solved analytically as provided by the Torsten library ready-to-use solver. Torsten package (Zhang et al., 2021) is based on Stan software (v2.27.0) and provides functions that facilitate the analysis of pharmacometrics data (Margossian et al., 2021). The library handles clinical event schedule data written in NONMEM (Beal et al., 2009) conventions and the computation of steady-state dosing^8^. A steady-state is reached when the quantity of drug eliminated matches the quantity of drug that reaches the systemic circulation. In the context of a repeat-dose regimen, steady-state concentrations of drugs in the blood will rise and fall according to a repeating pattern as long as the dosing schedule remains unchanged. Torsten also offers specific functions for one- and two-compartment models for which it implements analytical solutions with matrix exponential in Stan language.

## 3. Results

### 3.1. Bayesian inference on synthetic data

In order to validate the Bayesian workflow, a synthetic dataset is generated from a one-compartment population model with first-order absorption, based on the same dosing schedule as the clinical setting, and with population PK parameters as *CL*_*pop*_ = 7 (L/h), *V*_*pop*_ = 65 (L), 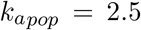 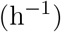. Inter-individual variations 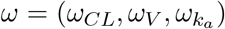 are equally set to 0.3, and residual intra-individual variation *σ* is set to 0.1. An example of simulated drug concentration data compared to empirical observation for a single individual (for 67-subject) is shown in Figure 1.

**Figure 1.**
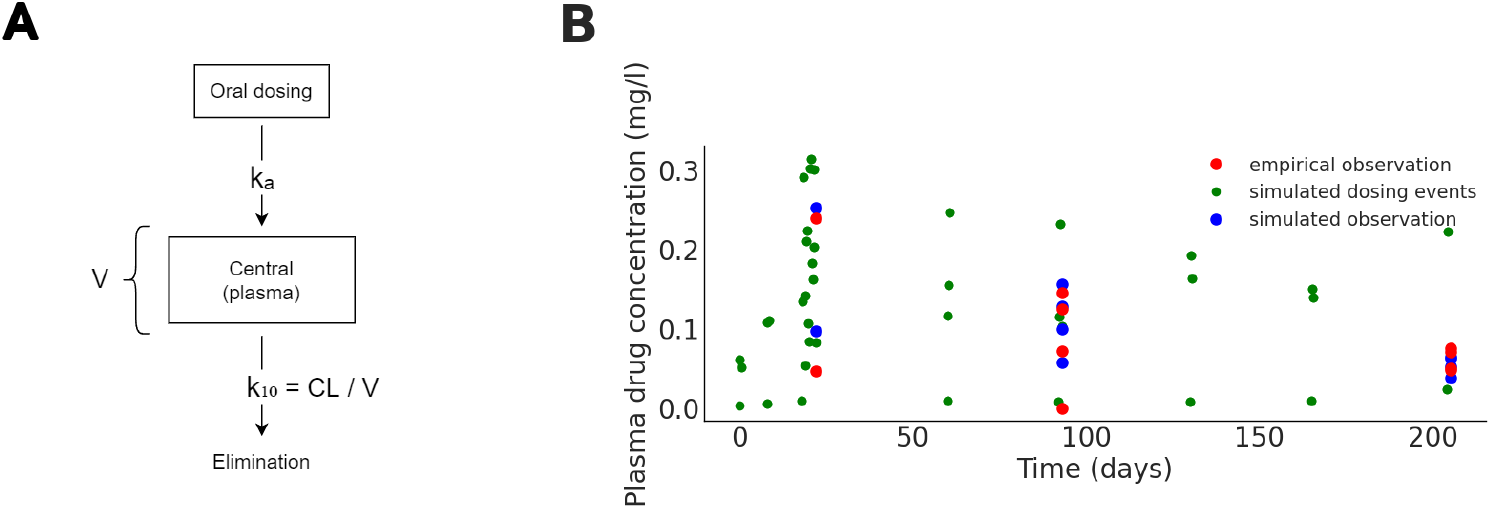
(**A**) Diagram of a one-compartment model; oral intake of drug is absorbed into the blood plasma (central compartment) with first order absorption rate *k*_*a*_ (h^−1^) and is eliminated from the plasma with constant *k*_10_ = *CL/V* (h^−1^). (**B**) One-compartment-simulated (in blue) and empirical (in red) data corresponding to plasma drug concentration (mg/L) for a single individual and additional simulated data (in green) from the same dosing schedule.

Bayesian inference of unknown parameters of the underlying data generating process depends on both observed responses and prior information about the underlying generative process. For the generated dataset, we aim to explore the actual sensitivity of the inference process to the level of information encoded in priors. To do so, we consider two different prior specifications on population parameters *CL*_*pop*_, *V*_*pop*_ and 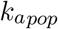, while prior distributions placed on other parameters remain identical. Inter-individual variations are modeled by a variance-covariance matrix Ω, in which for computational efficiency in Monte Carlo sampling, it is decomposed as

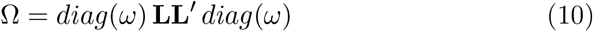

with **L** (resp. **L**′) as the Cholesky factor (resp. its transpose) of a correlation matrix *R* = **LL**′, and 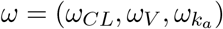 as the vector of standard deviation of PK parameters. Here, **L** is assigned with a Lewandowski-Kurowicka-Joe (LKJ; Lewandowski et al. (2009)) prior distribution (a distribution over the set of correlation matrices or their Cholesky factor) with uniform density over all Cholesky factors of correlation matrices, whereas *ω* is assigned with a half-normal prior with zero mean and standard deviation of 0.1:

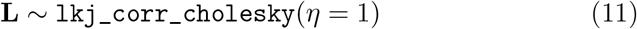

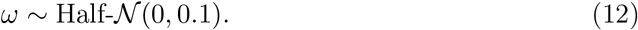

Individual parameters (normalised without covariate influence) denoted by 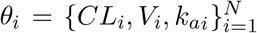 are modeled by a Gaussian distribution centered on population parameters 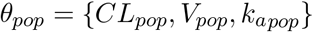,

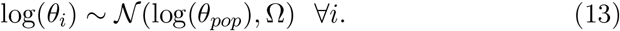

Given the parameters *θ*_*i*_ for the *i*^*th*^ individual, the virtual amount of drug in both absorption and plasma compartments is computed by analytically solving the one-compartment ODE system given by Eq. (1). Here, we employed a ready-to-use one-compartment model solver provided by Torsten package for event schedule specified data (Zhang et al., 2021). Estimation of drug concentration in the blood for individual *i*, with unknown *C*_*i*_ underlying observations *C*_*i*_[*obs*], is given by 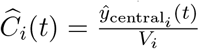, with the likelihood of

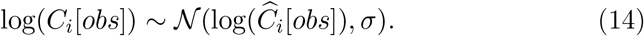

Given the above formulation, we compare two classes of *a priori* information on the population PK parameters. In the first set-up (denoted by model I), we place a weakly informative prior distribution centered around the true value of the parameters:

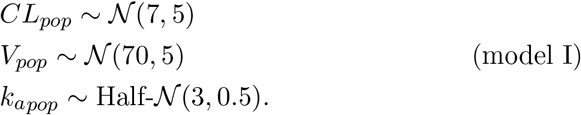

In the second set-up (denoted by model II), we place more diffuse (uninformative) priors but still centered on true value for parameters *V*_*pop*_ and 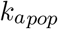:

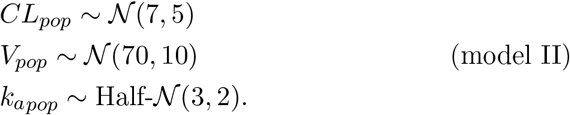

To perform inference, prediction, and model comparison, we ran four individual chains with NUTS default setting in Stan (e.g., 2000 discarded warm-up iterations and 2000 saved sampling iterations, expected acceptance probability of 0.85, and maximum tree-depth of 10). In the following, we first monitor the convergence of Markov chains, and then we show the estimated posterior distributions and posterior predictive checks for population PK parameters, and finally we demonstrate model comparison (by WAIC) to select the model that best explains the data.

#### Sampling diagnostics

Sampling properties of both Bayesian models with weakly informative and diffuse priors (given by model I and model II, respectively) are overall satisfactory (see Table 1). None of the models exhibited sampling pathologies such as divergent transitions or reaching maximum tree-depth. The values of the potential scale reduction factor are 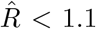 for all parameters in both Bayesian models, indicating the convergence of Markov chains to a stationary distribution. When considering a more exigent threshold, then 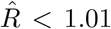 for all parameters of model I, except those relative to the 53^*th*^ individual. More precisely, the individual parameter which partly controls the individual variation for the 53^*th*^ individual and as a result biases the PK parameters and quantities (estimated and predicted concentration) for this individual. The Bayesian model II encounters more convergence issues with more individuals (16^*th*^, 44^*th*^, 53^*th*^, 64^*th*^). This model exhibits around 9% of its total 418 parameters that do not fall under the threshold against 2.1% for other model (see Table 1).

**Table 1.**
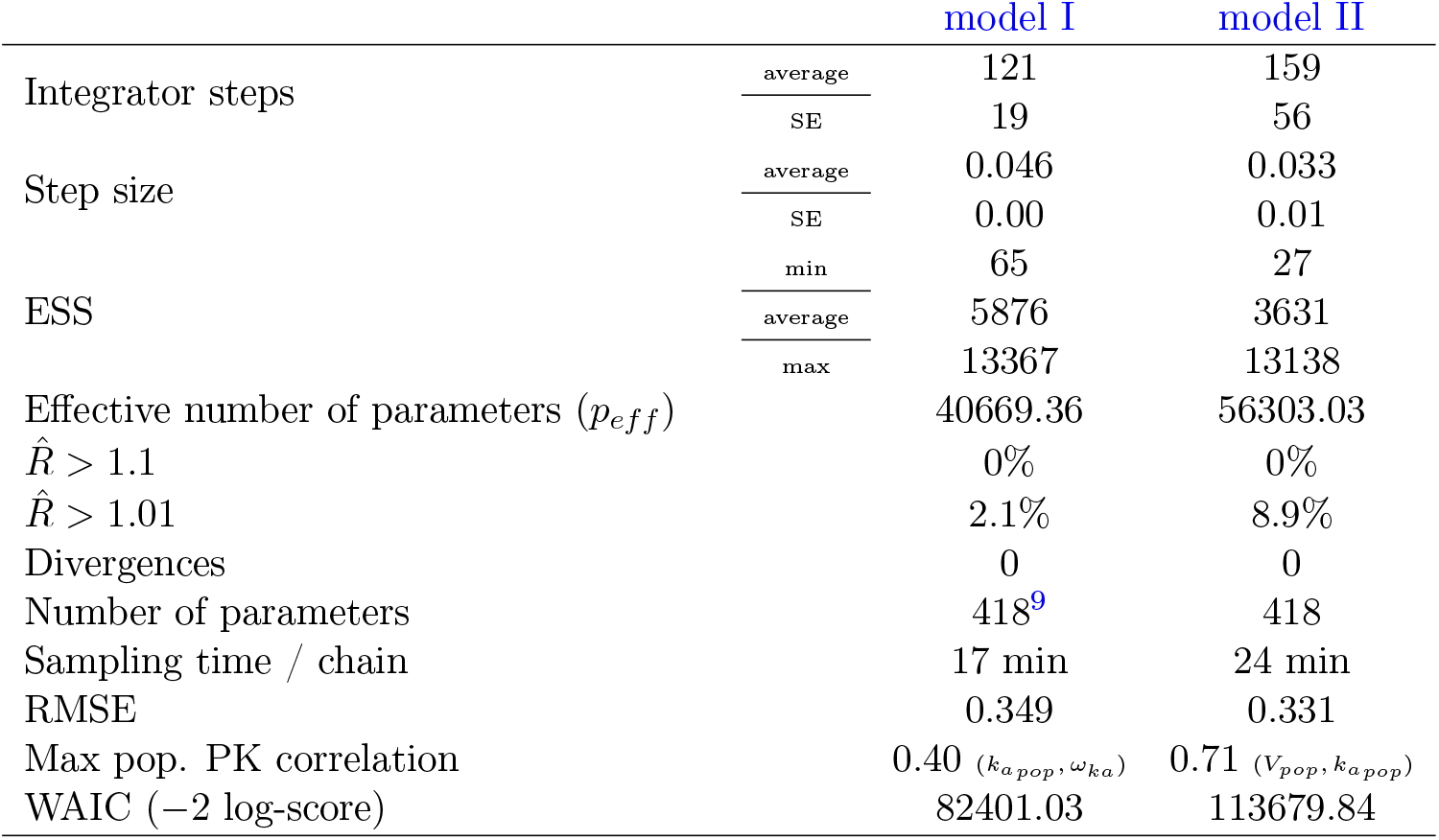
Summary statistics and convergence diagnostics for Bayesian inference on synthetic data (Figure 1), using weakly informative priors (given by model I) and diffuse priors (given by model II) averaged over four HMC chains. Overall, the Bayesian model I with a higher level of information in prior, yields faster computation, better convergence diagnostics, and substantially higher prediction accuracy.

The Bayesian model I also exhibits a better convergence according to HMC specific diagnostics compared to model II (see Table 1): the self-tuning step size of the leapfrog integrator is larger, and has fewer integration steps, resulting in faster computational time (around 1.5 order of magnitude), and more efficient sampling. Towards a principled Bayesian workflow (Gelman et al., 2020), we investigate more closely the convergence of the two fitted Bayesian models using the methods and quantities proposed by Vehtari et al. (2021) to address possible flaws in 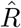-based convergence diagnostics.

#### Mixing of the chains

Rank plots are an alternative approach to trace plots for visual sanity checking of convergence in posterior chains (Vehtari et al., 2021) as shown in Figure 2. For comparison to raw trace plots of the chains for the population PK parameters see Appendix Figure B.10. Rank plots are histograms of posterior draws, ranked over all four chains, and plotted for each chain. If all chains are targeting the same posterior distribution, the rank histograms should not deviate significantly from uniformity, which is expected with a higher effective sample size. Figure 2 illustrates rank plots for population PK parameters using both Bayesian model I and model II. From this figure, we can see that a higher level of information in model I yields more uniform rank plots and thus better mixing of the chains.

**Figure 2.**
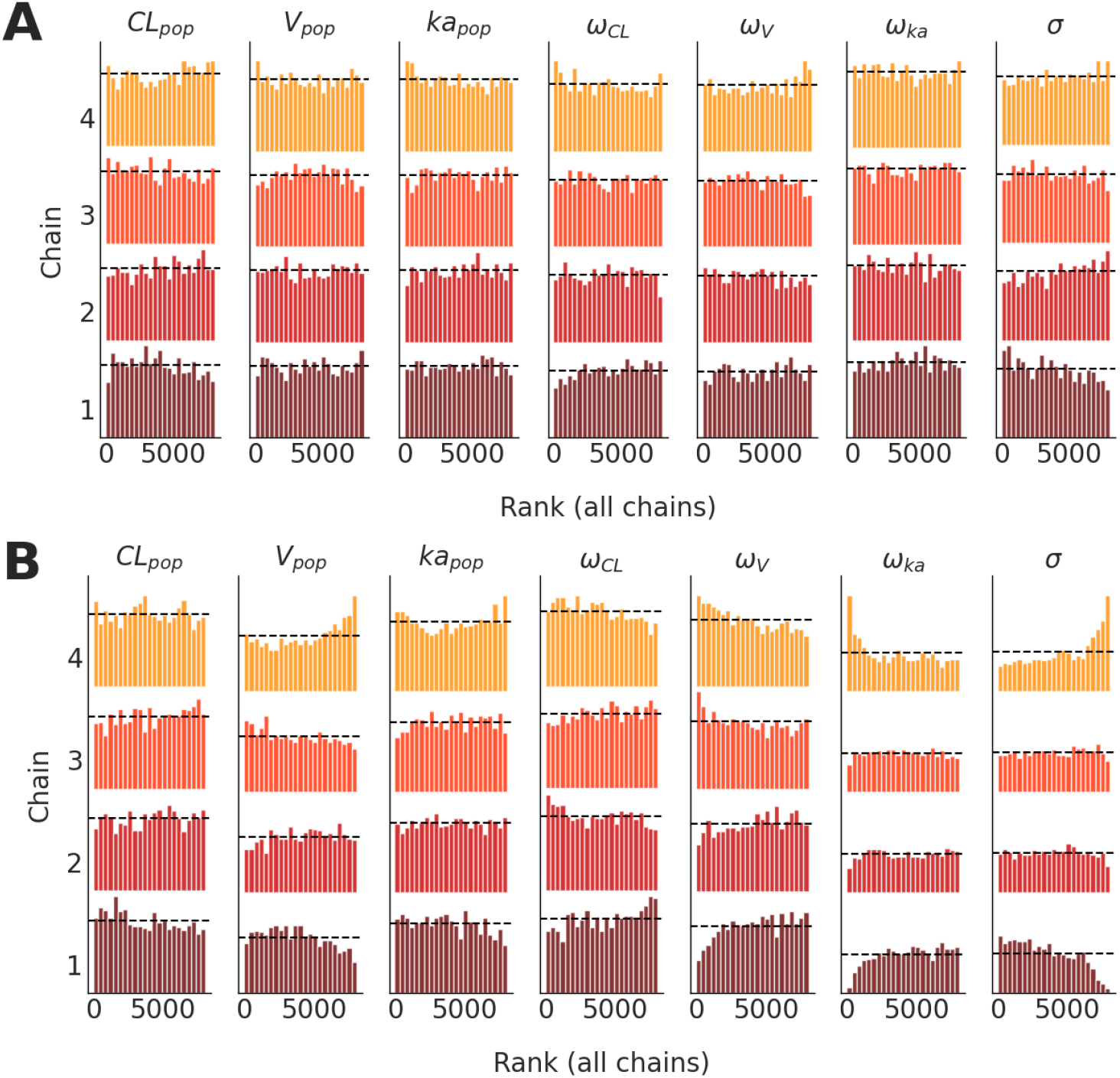
Rank plots (histograms of the 2000 ranked posterior draws over all four chains), for inference on the population PK and variability parameters using (**A**) Bayesian model I with weakly informative priors, and (**B**) Bayesian model II with diffuse priors. In the case of good mixing, all rank plots should be close to uniform (dashed horizontal lines). It can be seen that model I yields more uniform rank plots and thus better mixing of the chains.

#### Effective sample size

We have carefully monitored the effective sample size (ESS) as an indicator of the number of independent draws, which affects the uncertainty of the estimation. We investigate the ESS for the three population PK parameters using the computation of split-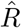 and ESS on rank-normalized (normal scores) samples, and corresponding diagnostics proposed by Vehtari et al. (2021). They have observed that the convergence of the chains is not necessarily uniform over the entire distribution of a parameter of interest, then proposed in addition to their rank-normalized based ESS (or bulk-ESS), a tail-ESS to evaluate the convergence in extreme quantiles.

In Figure 3, we illustrate the checks of efficiency of quantiles and small-interval probabilities, as well as the efficiency evolution with the number of iterations. The Bayesian model I yields an overall larger ESS for the three population PK parameters, especially for parameter *V*_*pop*_. The Bayesian model I quantile estimates efficiency is comparatively improved w.r.t. to model II for parameter *V*_*pop*_, and high quantiles of 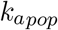, the first being higher and uniform and the latter being increased in the right tail of the distribution. The parameter *V*_*pop*_ in model II has noticeable small and decreasing efficiency in high quantiles. *CL*_*pop*_ quantile estimates efficiency is similar in both models. In the case of a well-behaved sampling, the estimated effective sample sizes should increase linearly with the number of iterations. This is the case for the population PK parameters in model I. However, in model II, the parameter *V*_*pop*_ exhibits a problematic drop at half of the total iterations in tail-ESS, which indicates inter-chain scale discrepancy. Population PK parameters of both models reach an acceptable ESS over the recommended threshold of 400 when the number of draws exceeds 2000, except for pathological parameter *V*_*pop*_ in model II. Together, these diagnostics validate that the samples in model I have better convergence to the target distribution.

**Figure 3.**
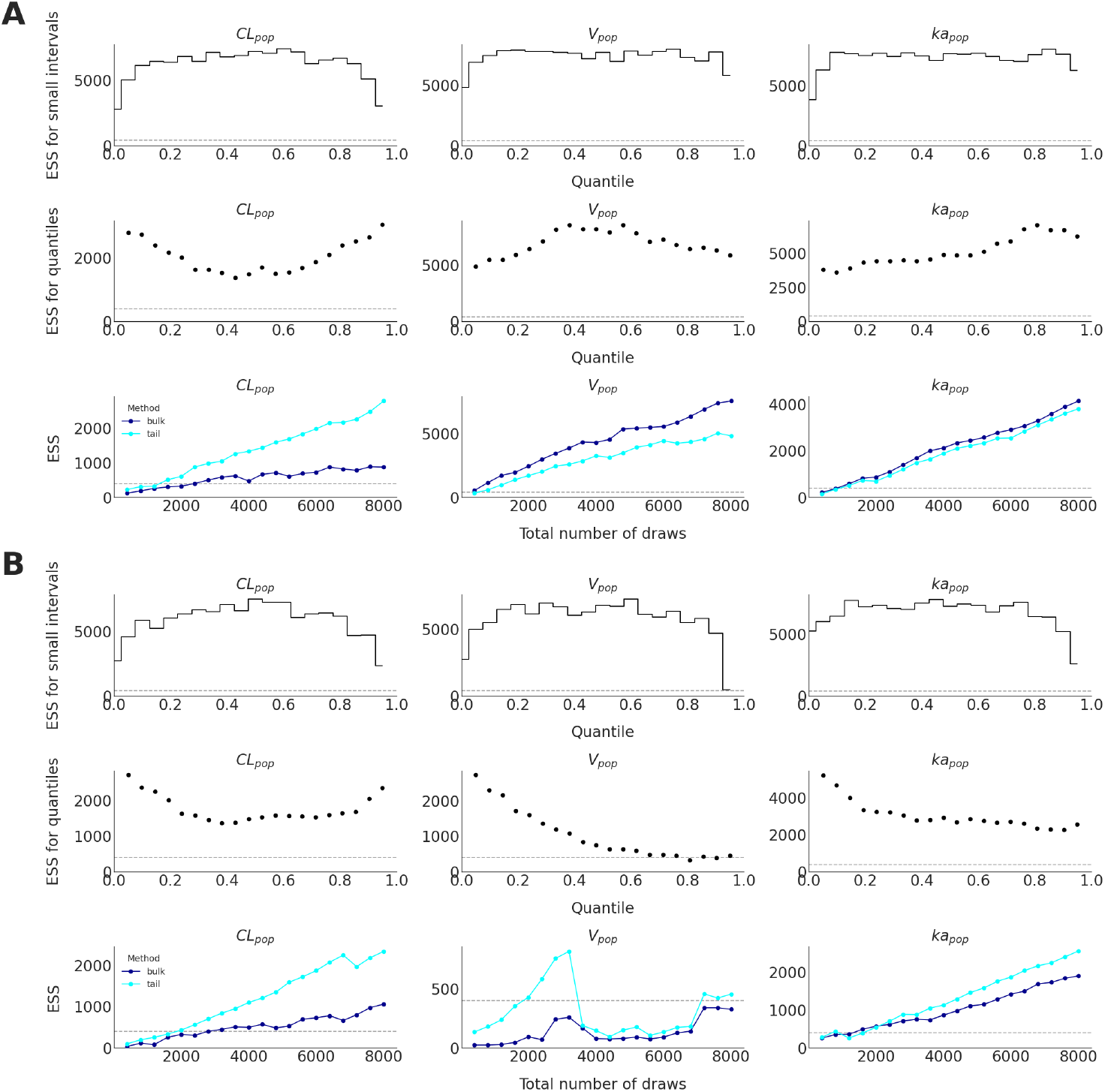
Effective sample size (ESS) plots of population PK parameters using (**A**) Bayesian model I with weakly informative priors, and (**B**) Bayesian model II with diffuse priors. In each panel, Top: local efficiency of small interval probability estimates, Middle: efficiency of quantile estimates, Bottom: estimated effective sample size (bulk and tail methods) with an increasing number of iterations. Bulk effective sample size: between-chains discrepancy location; Tail effective sample size: between-chains scale discrepancy. In the case of a well-behaved sampling, the estimated effective sample sizes should increase linearly with the number of iterations. Overall, a higher level of information in model I yields larger ESS, i.e., more independent posterior draws, thus, the more reliable estimations.

#### Posterior behaviour

The prior and posterior distributions of model I and model II are illustrated in Figure 4. It can be seen that taking model I, the true values of all PK parameters (dashed vertical lines) are well under the support of the posterior densities, indicating that the Bayesian parameter recovery was successful. In contrast, the setting in model II failed to recover the parameters *V*_*pop*_ and 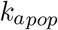. When using a wider set of priors (model II), the posterior distribution of *V*_*pop*_ shifts towards values that are significantly lower than ground-truth under over-influence of the data (over-fitting). A similar phenomenon is observed for the absorption parameter *k*_*a*_. The uncertainty of the estimation of posterior quantities (mean, MCSE of the mean, standard deviation, 95% credible intervals, and ESS) for the population PK parameters in both Bayesian models are summarised in Table 2.

**Figure 4.**
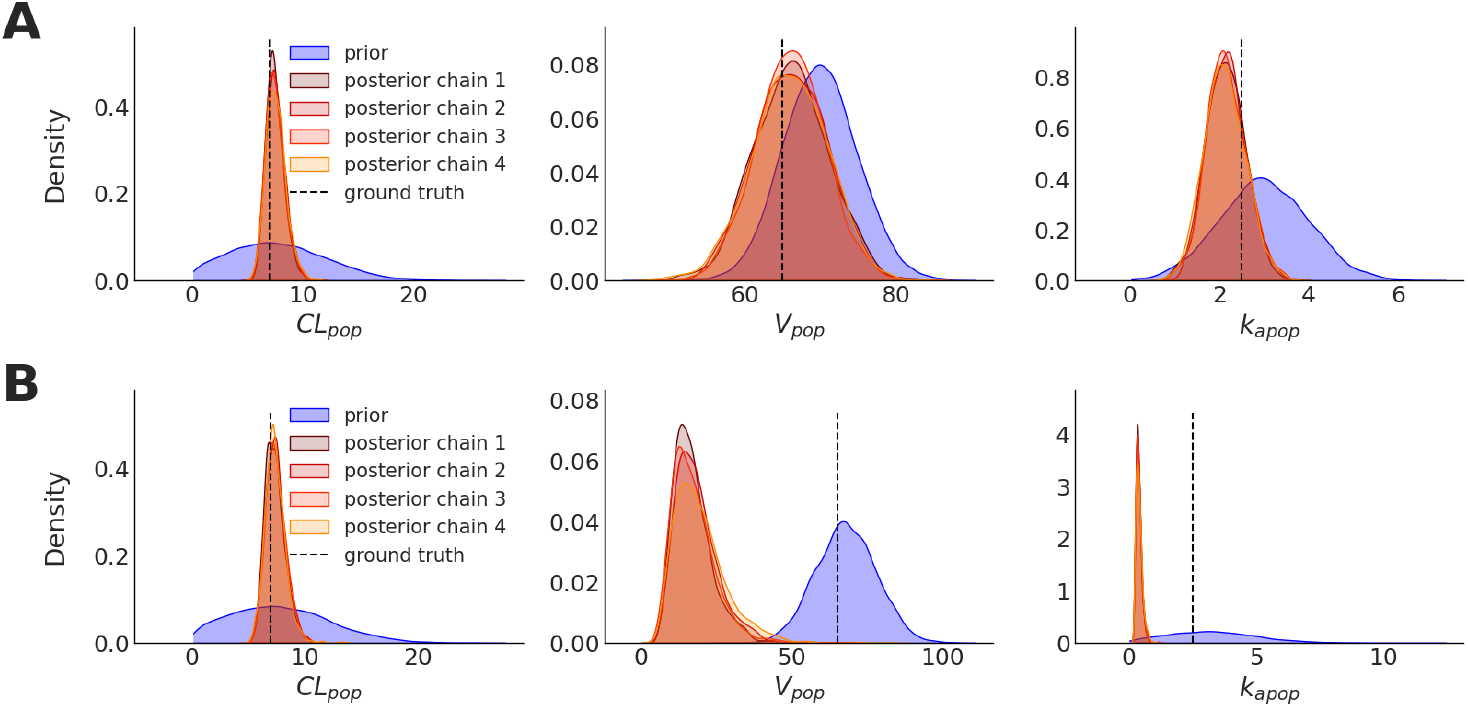
Comparison of prior and posterior distributions (4 chains, 2000 draws each) of population PK parameters for the two models fitted against simulated data: (**A**) Bayesian model I with weakly informative priors, and (**B**) Bayesian model II with diffuse priors. The dashed vertical lines indicate the true values.

**Table 2.**
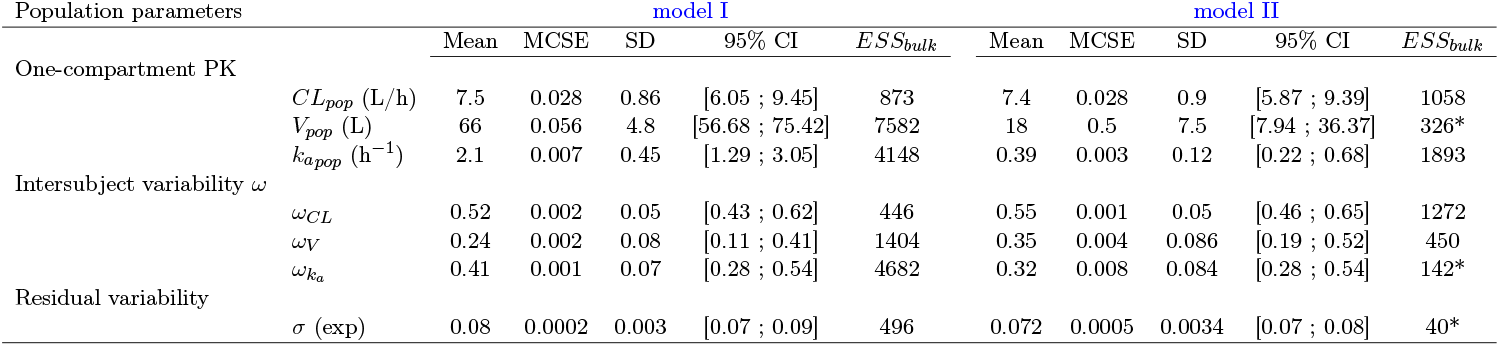
Comparison of sampled posterior distributions of population PK parameters with Bayesian model I and model II; sample mean, MCSE, sample standard deviation (SD), posterior 95% credible intervals and effective sample size (*ESS*_*bulk*_). Parameters marked with a * have ESS below the limit of 400 recommended by Vehtari et al. (2021).

Visualizing the joint posterior samples in pair plots is especially useful for identifying collinearity between parameters as well as the presence of non-identifiability (banana-like shapes). As shown in Figure 5, model II exhibits a larger degenerate behavior in the form of a positive correlation (*r* > 0.7) between *V*_*pop*_ and 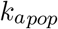 posterior samples that is not seen in the more informative model I (*r* < 0.2). Such a high collinearity leads to an inefficient exploration of the posterior which can be quantifiably observed in decreased numbers of effective samples and increased 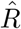 values. Moreover, the estimation of posterior density of *CL*_*pop*_ is more robust than the other two parameters. The robustness of the sampling of *CL*_*pop*_ w.r.t. other parameters was also observed in the ESS plots (Figure 3). According to these results, the Bayesian set-up used in model I substantially improved the convergence of the recovery for population PK parameters.

**Figure 5.**
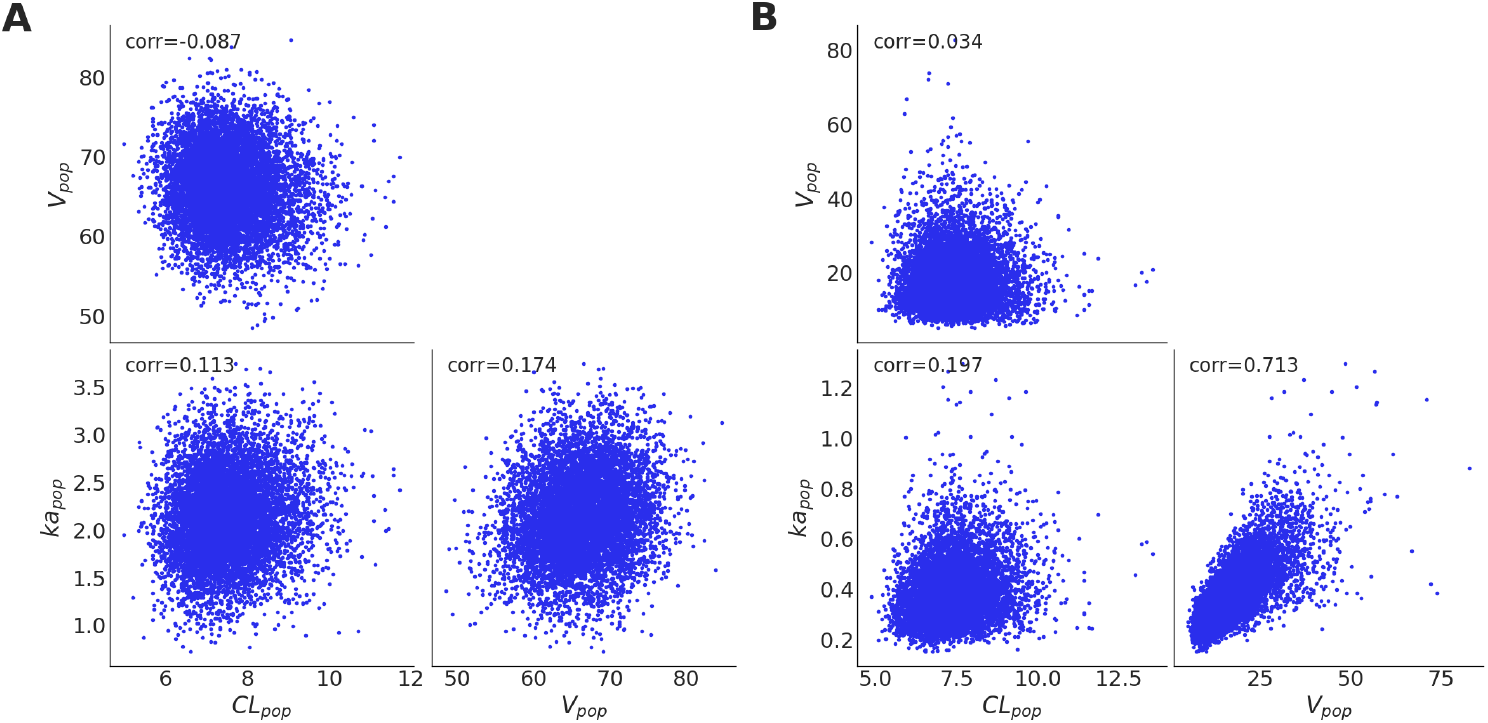
Paired posterior draws (4 chains, 2000 draws each) of population PK parameters using (**A**) Bayesian model I with weakly informative priors, and (**B**) Bayesian model II with diffuse priors. In model I, joint posteriors are well decorrelated (independent samples), whereas in model II, joint plot of *V*_*pop*_ and 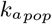 exhibits high positive correlation (sample Pearson correlation coefficient *r* = 0.71).

#### Posterior predictive checks

Posterior predictive (or retrodictive) checks are a widely used tool to verify the reliability of the fitted model by monitoring its predictive performance on the fitted dataset. Synthetic datasets (67 individuals, total of 427 observations) of plasma drug concentration are repeatedly drawn from the predictive distribution given by the fitted model (4 chains, each with 2000 draws). Punctual predictions (mean) and the corresponding 95% credible intervals are extracted from the draws and plotted along with the actual observations (see Figure 6). For both Bayesian model I and model II, these visual checks suggest no pathological bias in the predictive distribution compared to the fitted data.

**Figure 6.**
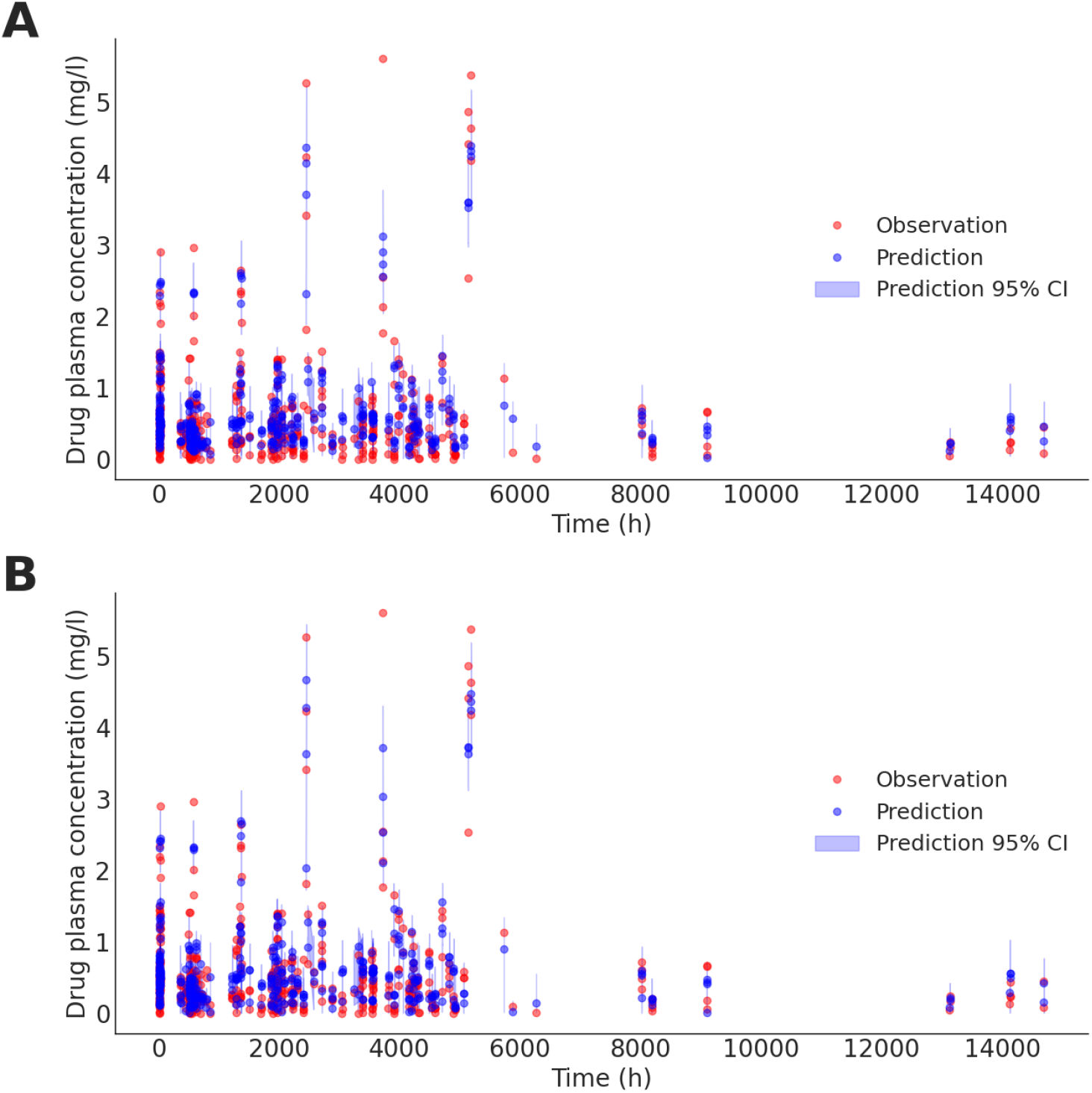
Posterior predictive checks of individual drug plasma concentration (mg/l) - all individuals plotted (in red), using (**A**) Bayesian model I with weakly informative priors, and (**B**) Bayesian model II with diffuse priors. Point prediction (mean, dark blue) is plotted along with its 95% CI (in light blue). If successive observation records are spaced by less than 7 days, the corresponding posterior 95% CI is plotted as a shaded area; else as an error bar.

#### Residuals

Non-normalised residuals are calculated as differences between observed plasma concentration *C*_*obs*_ and the estimated concentration computed by solving the one-compartment ODE system (see Eq. (1)), over the fitted individual PK parameters 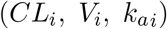 and evaluated at observation points 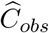. Monitoring the residuals can help in pointing out issues in the estimation of pharmacokinetic parameters, an inadequate compartmental modeling, or ODE-solver inaccuracies. Histograms of non-normalised residuals for both model I and model II are given in Appendix Figure B.11. They are reasonably zero-centered (not biased), and there is no significant difference in distributions of non-normalised residuals of model I and model II. Note that the root mean squared error (RMSE, Figure B.11, and Table 1) as a within-sample predictive accuracy is slightly smaller in model II; this is consistent with the marginal posterior densities (Figure 4) where the interaction of the likelihood with more diffuse priors fails to recover the true parameters; hence, the resulting model fits the observed data even better but at the cost of a poor representation of the true data generating process (Betancourt, 2018; Gelman et al., 2017).

#### Model selection

Lastly, to select the best Bayesian model considered in this study for the synthetic dataset, we use the widely applicable information criterion (WAIC) as a measure of model evidence. WAIC is an indicator for the comparison of point-wise out-of-sample predictive accuracy of Bayesian models, based on the whole estimated posterior distribution. WAIC is here reported (see Table 1) in the −2 log-score scale (deviance): a model with a smaller WAIC value suggests a higher predictive accuracy, i.e., a better balance between accuracy and complexity, though the number of parameters is equal for both candidate models. Here, the model comparison based on WAIC substantially favors model I (since ΔWAIC ≫ 10). Taken together, the Bayesian set-up used in model I substantially improved the convergence and model predictive power for estimation over population PK parameters.

### 3.2. Bayesian inference on clinical data

After calibrating models on simulated data, we fit model I on empirical Baclofen data. In particular, we use the set of population PK priors defined in model I. The same as previous section, we ran 4 chains of 2000 iterations each, with default settings in Stan (e.g., 2000 warm-up iterations, target Metropolis acceptance rate of 0.85, and a maximum tree-depth of 10).

#### Sampling diagnostics

Stan diagnostics reported no warning on convergence, in particular, there is no 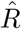 value above the threshold of 1.01, no diverging transition, and no quantity with insufficient ESS. We also rely on convergence diagnostics as shown in the investigation of models on simulated data: trace plots in Figure C.12 show that the chains are mixing well; rank plots in Figure C.13 are close to uniform; effective sample size (bulk and tail methods) in Figure C.14 is large for the main three population PK parameters; paired samples of population PK parameters in Figure C.15 exhibit no significant correlation.

#### Posterior distributions

The sampled posterior distributions of the three main population PK parameters and their respective inter-individual variability parameters are shown and compared to their prior distributions in Figure 7. Statistical summary of posterior distributions of the population PK parameters is summarised in Table 3. We report the mean, MCSE of the mean, standard deviation, 95% credible intervals, and ESS. Interpretation of Bayesian posterior credible intervals is straightforward: a 95% CI is the central portion of the posterior that concentrates 95% of sampled values; given the observed data, the parameter falls into the interval with probability of 95%. Distributions of sampled individual PK parameters *CL*_*i*_, *V*_*i*_ and 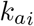 are plotted for every individual in Figure C.16.

**Figure 7.**
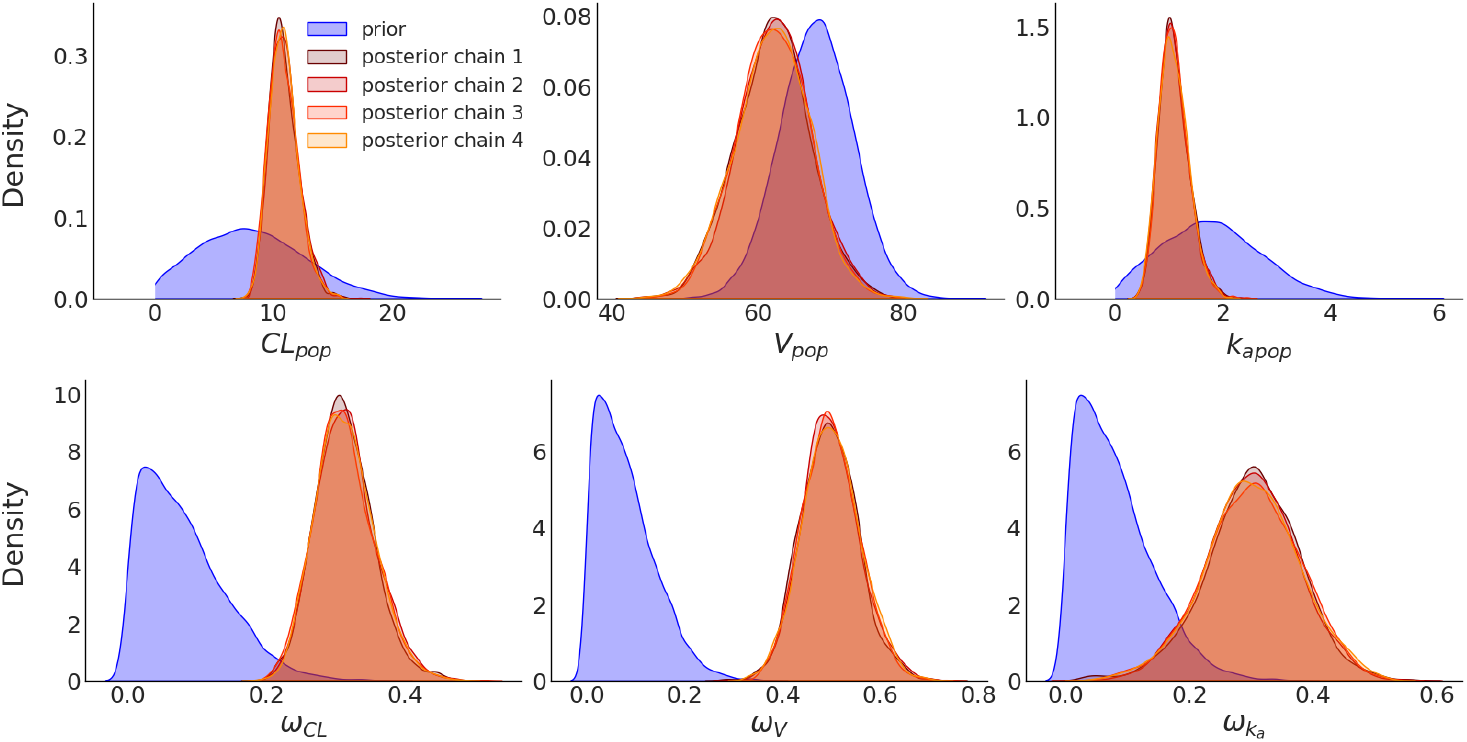
Comparison of prior and posterior distributions (4 chains, 2000 draws each) of population PK parameters and their respective variability parameters for weakly informative model I, fitted against empirical Baclofen data.

**Table 3.** Posterior sampled distributions of population PK parameters from empirical Baclofen data; sample mean, Monte-Carlo standard error (MCSE), sample standard deviation (SD), and posterior 95% credible interval.

| Population parameters |  | Posterior sampled distributions |  |  |  |  |
| --- | --- | --- | --- | --- | --- | --- |
| | | Mean | MCSE | SD | 95% C.I. | $ESS_{bulk}$ |
| One-compartment PK |  |  |  |  |  |  |
| | $CL_{pop}$ (L/h) | 11 | 0.023 | 1.2 | [8.80 ; 13.68] | 2861 |
| | $V_{pop}$ (L) | 62 | 0.05 | 5.1 | [52 ; 72.07] | 10419 |
| | $k_{a_{pop}}$ (h<sup>-1</sup>) | 1.1 | 0.0041 | 0.27 | [0.64 ; 1.72] | 4307 |
| Intersubject variability $\omega$ | | | | | | |
| | $\omega_{CL}$ | 0.31 | 0.0006 | 0.042 | [0.24 ; 0.40] | 4421 |
| | $\omega_V$ | 0.5 | 0.0008 | 0.058 | [0.39 ; 0.62] | 5400 |
| | $\omega_{ka}$ | 0.3 | 0.0013 | 0.076 | [0.15 ; 0.45] | 3354 |
| Residual variability |  |  |  |  |  |  |
| | $\sigma$ (exp) | 0.084 | 0.00003 | 0.003 | [0.08 ; 0.09] | 10805 |

#### Posterior predictive checks

Predicted Baclofen plasma concentration datasets (*N* = 4 chains × 2000 draws) are generated from the estimated predictive distribution (see Figure 8). Each shown data-point corresponds to an actual individual observation (in red). The posterior predictive checks are shown for all individual observations (in blue). For every individual observation, the corresponding punctual prediction (mean) and 95% confidence interval over the repeated simulations are also plotted (in light blue). No systematic deviation of predictive simulations from reality is visible, suggesting accurate capturing of plasma concentration by the fitted model’s predictive distribution.

**Figure 8.**
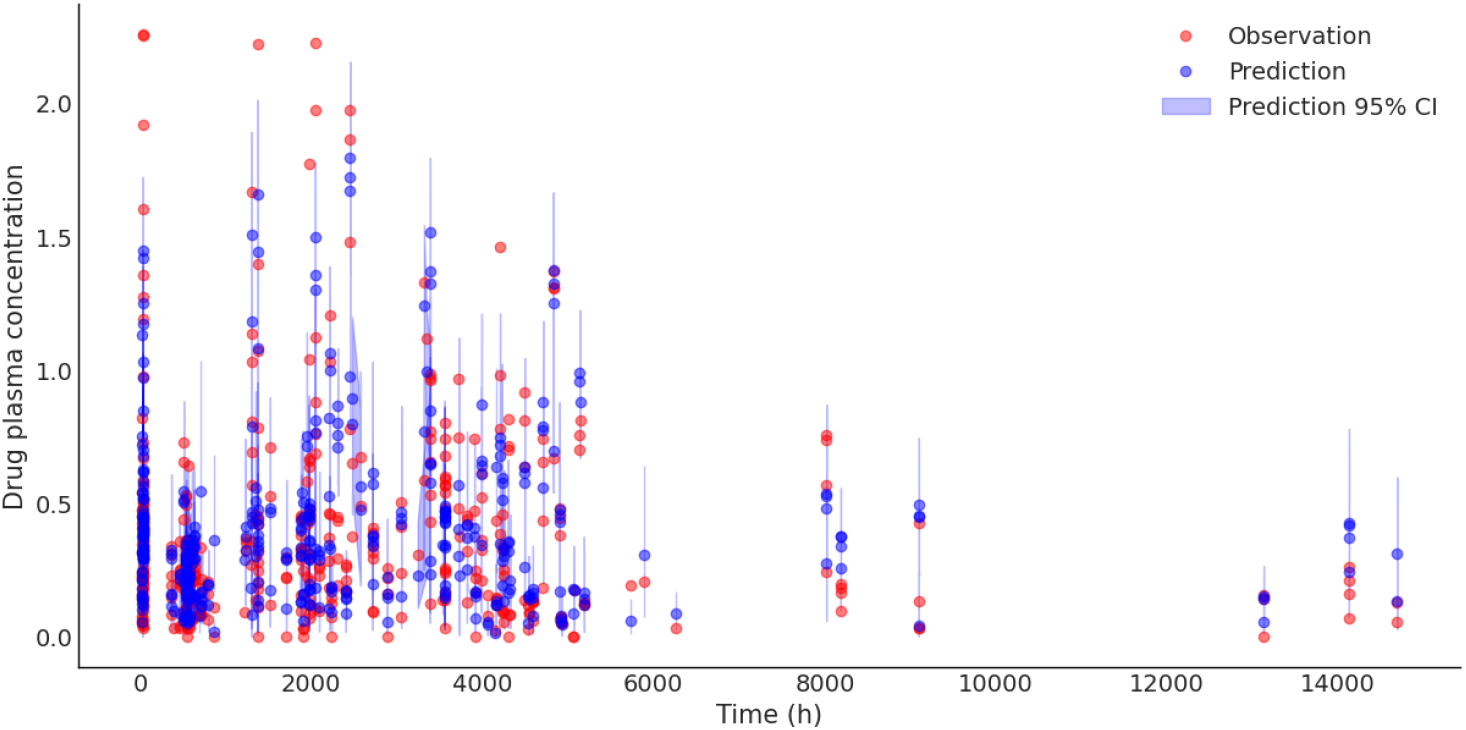
Posterior predictive checks (in dark blue) of individual concentration of Baclofen in blood plasma (mg/l) using model I, fitted against empirical data (in red) for all individuals. Point prediction (mean, in dark blue) is plotted along with its 95% CI (in light blue). If successive observation records are spaced by less than 7 days, corresponding posterior 95% CI are plotted as a shaded area; else as an error bar.

#### Residuals

Checking of residuals between the fitted ODE-solved estimated concentration at observation points 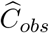 and actual observations *C*_*obs*_ allows ensuring that there is no bias in the estimation of the PK parameters and issue in resolution of the underlying ODE system. Histogram of non-normalized residuals and the residuals along the time-axis are shown in Figure 9, indicating no biased estimation.

**Figure 9.**
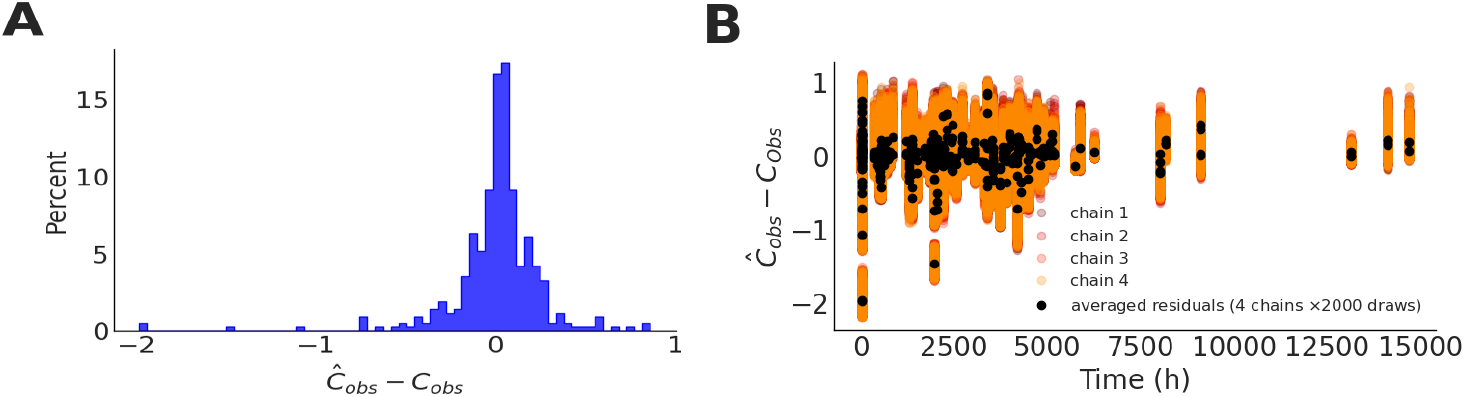
Residuals (non-normalised) plots of fitted ODE-solved estimated concentration at observation points 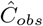 minus actual observations *C*_*obs*_. (**A**) Residuals histogram, and (**B**) Residuals as a function of time.

#### Covariates

Clinical data includes 18 biological covariates which are detailed in Imbert et al. (2015). The influence of a continuous covariate (denoted by COVAR) on individual parameter *θ*_*i*_ is modeled by

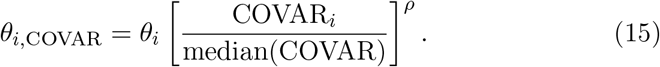

Searching for significant covariates on PK parameters was performed by evaluating repeated regressions (in linear and nonlinear forms of Eq. (15)) on HMC-fitted individual PK parameters. The statistical significance (*p* < 0.01) of covariate coefficients such as Gender, Body weight, Age, Height, Creatinine serum, Creatinine clearance, Urea, Alanine aminotransferase, Aspartate aminotransferase, Albumin, Mean corpuscular volume, Prothrombin ratio, Fibrinogen, Gamma-glutamyl transferase, Alkaline phosphatase, Carbohydrate-deficient transferrin, Tobacco, and Fagerstrom score, was tested for each fit. None exhibited a significant difference for more than half of the fits. As a result, no covariate was retained. This is consistent with the covariate selection that was performed with a NONMEM (Beal et al., 2009) routine in Imbert et al. (2015) and resulted in no selected covariate.

## 4. Discussion

Bayesian inference is a principled method to estimate the posterior distribution of unknown quantities given only observed responses and prior beliefs about unobserved hidden states (or latent variables). An advantage of using the Bayesian framework in the context of inference/prediction is the ability to generate not only a single point estimate (e.g., in the Frequentist approach), but also full probability distributions for the quantities of interest (uncertainty quantification for decision-making process). From the latter, one can directly extract quantiles, with the possibility to answer questions such as "what is the probability that the parameter of interest is greater/smaller than a specific value?", with the confidence intervals in estimation. In addition, the propagation of uncertainty in the Bayesian framework provides a more robust and reliable predictive capability of the model under study, rather than point estimation with optimization methods. In particular, the out-of-sample prediction accuracy (i.e., the measure of the model’s ability in new data prediction e.g., using WAIC) enables reliable and efficient evaluation of potential hypotheses, as performed in this study. Several previous studies have used a scoring function (such as root mean square error or correlation) to measure the similarity between empirical and fitted data (Imbert et al., 2015). The choice of scoring function can dramatically affect the ranking of model candidates, and ultimately the decision-making processes (see RMSE in Table 1). Rather, we used non-parametric probabilistic methodology to analyze data, while various convergence diagnostics were monitored to assess when the sampling procedure has converged to sampling from the target distribution.

Another advantage of the Bayesian approach used in this study is to integrate our prior information (domain expertise before seeing the data) in the inference process to better improve the model’s ability in prediction of new data. For instance, it has been shown that measuring the prediction accuracy of a whole-brain network model with a higher level of information in priors provides decisive evidence in favor of the true hypothesis regarding the degree of epileptogenicity across different brain regions (Hashemi et al., 2021; Vattikonda et al., 2021). Accuracy of the predictive models is critical to determine the quality of their predictions for evidence-informed decision-making. It is important to note that the prior information is relevant in estimating the parameters rather than the calculation of predictive accuracy (Gelman et al., 2014b). However, a substantial change in priors will affect the computations of the marginal likelihood, thus, the accuracy of the predictive models. An inappropriate choice of prior can lead to weak inferences and thus poor predictions (see Figure 4), whereas a sufficient amount of information encoded through the prior distribution can provide decisive evidence in favor of the correct hypothesis, as shown in this study (see WAIC in Table 1). In addition to achieving a model with higher performance prediction accuracy, the informative priors can dramatically improve the exploration of the search space in terms of computational cost and algorithmic diagnostics such as effective numbers of samples, rendering the inference more efficient (see Table 1, Figure 2, Figure 3, and Figure 5).

Monte Carlo methods allow us to sample from and hence, approximate the exact posterior densities, in which the sampling process does not require knowledge of the whole distribution. Hamiltonian Monte Carlo (HMC) uses the gradient information of the posterior to avoid the undesired random walk of the traditional sampling algorithms, thereby samples efficiently from posterior distributions with correlated parameters, in particular in high dimensional search spaces. Thanks to open-source Stan programs (Carpenter et al., 2017), the automatic HMC sampling refined the inference on population PK parameters with the uncertainty in estimated parameters compared to point estimation in previous work of Imbert et al. (2015). In this study, we integrated the domain expertise in inference through the prior distribution to obtain the full posterior distribution that is well consistent with the data and domain expertise e.g., prior information refined inference for the clearance parameter *CL*_*pop*_ and its variability *ω*_*CL*_ that are outside the 95% bootstrap confidence intervals compared to previous analysis. The posterior distributions of other inferred population PK parameters exhibit mean values that are in the previously proposed confidence intervals, and the credibility intervals extracted from their posterior distributions have been narrowed in comparison. Moreover, here the residual variance is reduced by a factor of 10.

The Bayesian approach relying on advanced MCMC sampling algorithms used in this study provided accurate and reliable estimation of Baclofen effect, validated by posterior behavior analysis and various convergence diagnostics. Such a principled and probabilistic methodology enabled us to integrate the prior information in the explanation of observations to maximize the model prediction power for a given patient with AUD. This work offers proper guidance for the prediction of drug efficacy in clinical practice to bring personalized medicine closer to reality in the treatment of brain disorders.

## Footnotes

1 For oral dosing, doses are added in the dosing compartment; infusion and IV bolus injections must be added directly in the central compartment.

2 Matrix exponential *e*^*t***A**^ is defined as 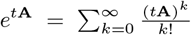. Stan features a matrix exponential function `matrix_exp`.

3 Log predictive density or log-likelihood is proportional to the mean squared error if the model is normal with constant variance.

4 The effective number of parameters is calculated by summing over all the posterior variance of the log predictive density for each data point.

5 *W* is defined as the averaged sum of the squares of within-chain variance of every chain (Gelman et al., 2014a).

6 A simple example: two chains with complementary increasing and decreasing values will yield 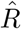 close to 1, even though they are not mixing.

7 A numerical integration scheme for Hamiltonian systems to nearly conserve the total energy and is particularly useful when treating long times.

8 Computation of steady-state dosing in Torsten is detailed in Appendix D in the case of one-compartment modeling.

9 Three population parameters 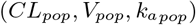 and their respective variability 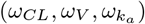; 3*×*67 individual parameters 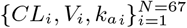; elements of 3×3 Cholesky matrix **L**; 3 × 67 standard normal variables for non-centered parametrization of Eq. (13); residual variability *σ*.

10 (https://github.com/metrumresearchgroup/Torsten)

## Information Sharing Statement

The patient datasets cannot be made publicly available due to the data protection concerns regarding potentially identifying and sensitive patient information. Interested researchers may access the datasets by contacting Nicolas Simon at Aix-Marseille Université. The main source codes needed to reproduce the presented results are available on GitHub (https://github.com/ins-amu/Stan_PKPD_examples).

## Acknowledgements

This work was funded by Human Brain Project SGA3, and the Fondation pour la Recherche Médicale (DIC20161236442) to V.K.J. The funders had no role in study design, data collection and analysis, decision to publish, or preparation of the manuscript.

## Author contributions

M.H., N.S, and V.K.J. designed the study. N.S acquired the data. N.B. performed the analysis. N.B., M.H, N.S, and V.K.J. wrote the manuscript.

## A. Hamiltonian equations of motion

## A.1. Expression of the Hamiltonian

The Hamiltonian function is expressed in terms of parameter *θ* and sampled auxiliary momentum *ρ* by developing the joint probability formula,

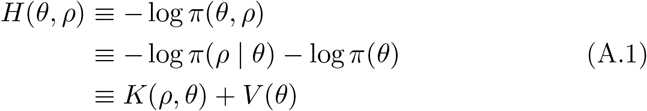

where *K*(*ρ*, *θ*) corresponds to the kinetic energy and *V* (*θ*) is the potential energy.

## A.2. Hamiltonian equations of motion

Parameters (*θ*, *ρ*) in phase space are evolved through Hamiltonian dynamics by solving the Hamiltonian equations of motion:

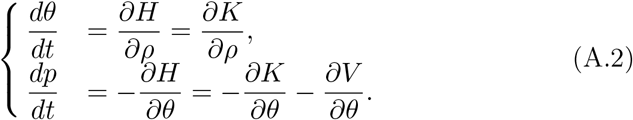

## A.3. Leapfrog integrator

For a number of iterations *L*, the leapfrog integrator successively updates the momentum of half a *ϵ* step then parameters of a complete step and then updates the momentum of half a step again.

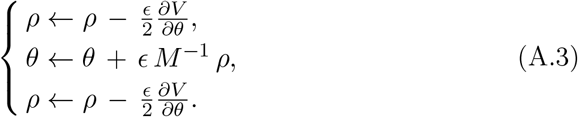

## B. Bayesian inference on synthetic data

**Figure B.10.**
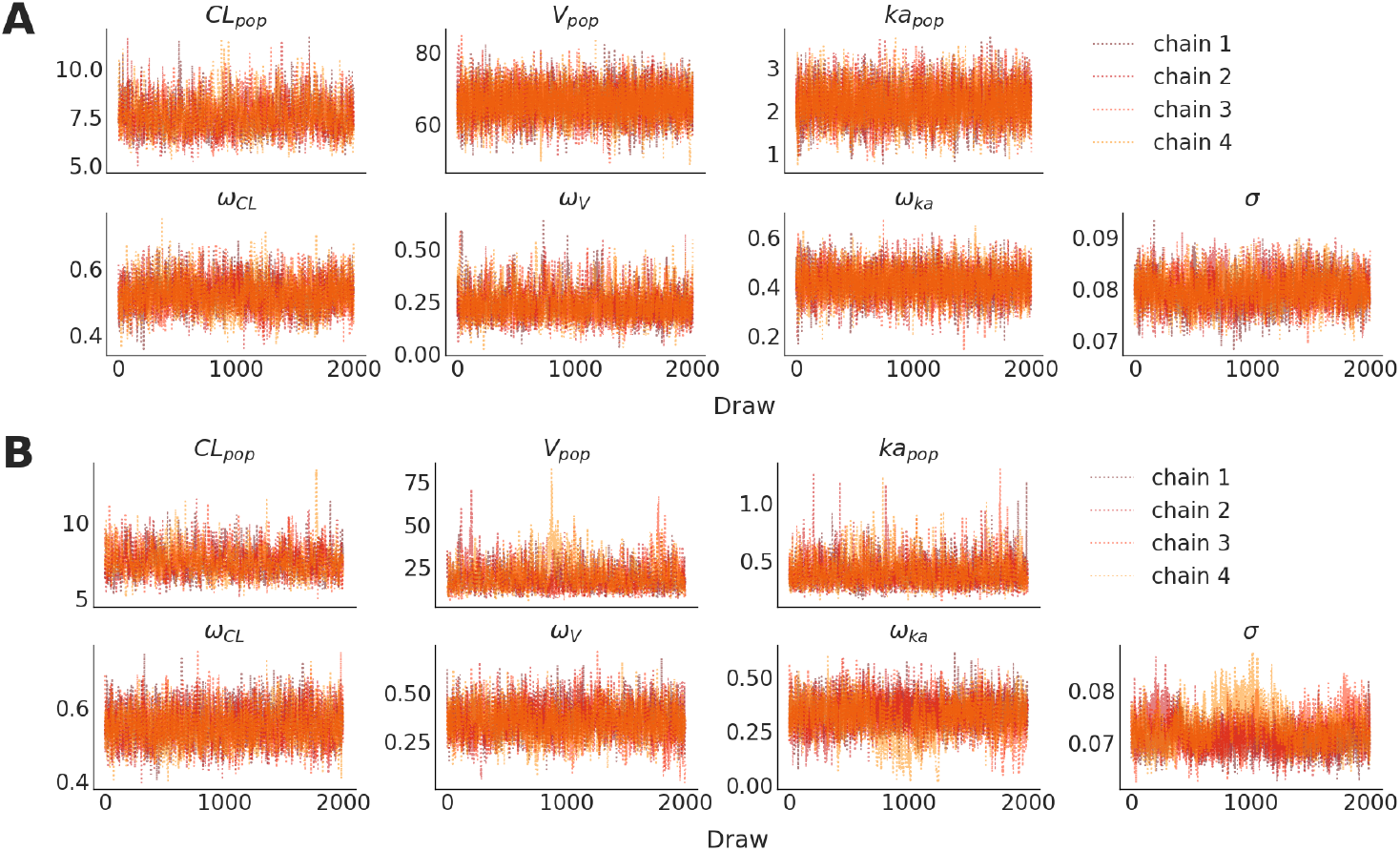
Trace plots of sampled posteriors for population PK parameters (4 chains, 2000 draws each) on synthetic data, using (**A**) Bayesian model I with weakly informative priors, and (**B**) Bayesian model II with diffuse priors.

**Figure B.11.**
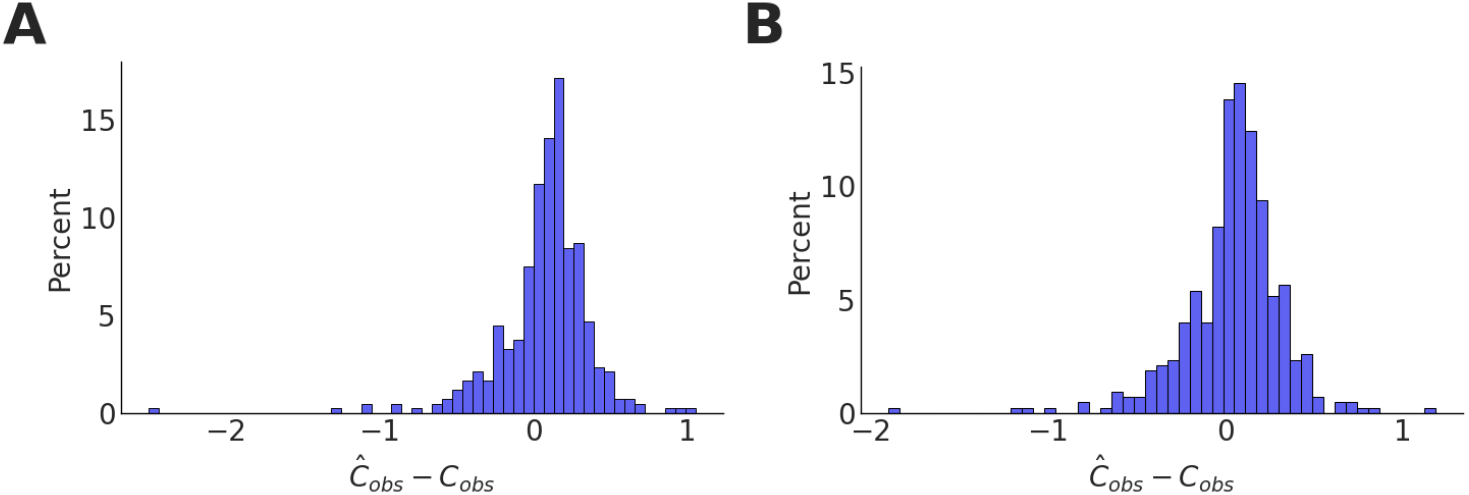
Histogram of residuals (non-normalised) for (**A**) Bayesian model I with weakly informative priors (RMSE = 0.348), and (**B**) Bayesian model II with diffuse priors (RMSE = 0.331).

## C. Bayesian inference on empirical data

**Figure C.12.**
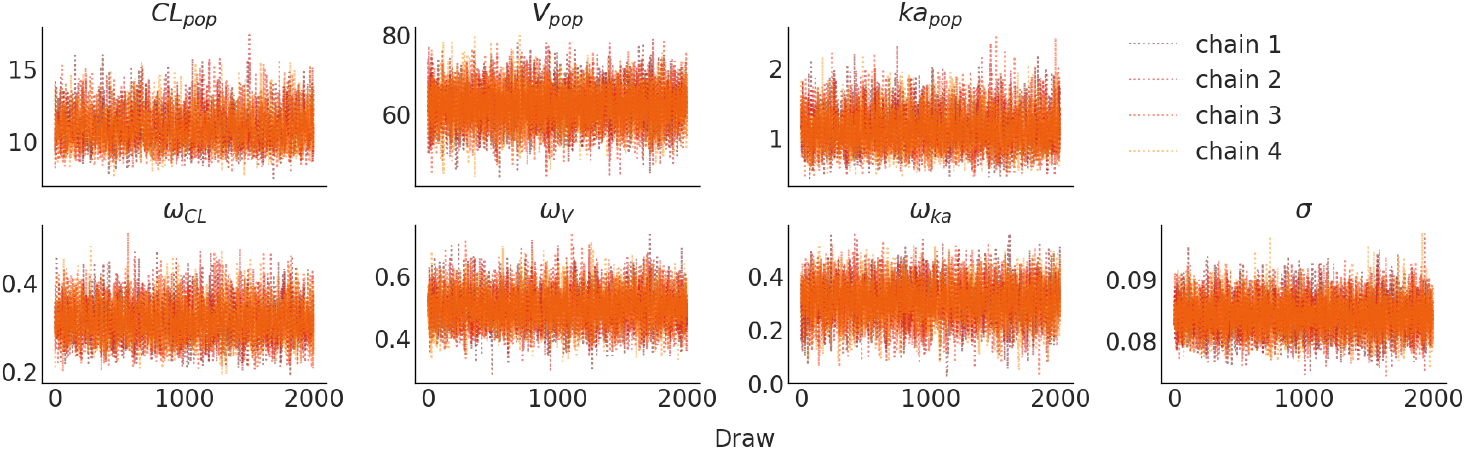
Posterior chains of population PK parameters (4 chains, 2000 draws each) for weakly informative model I fitted against empirical Baclofen data show that the chains are mixing well.

**Figure C.13.**
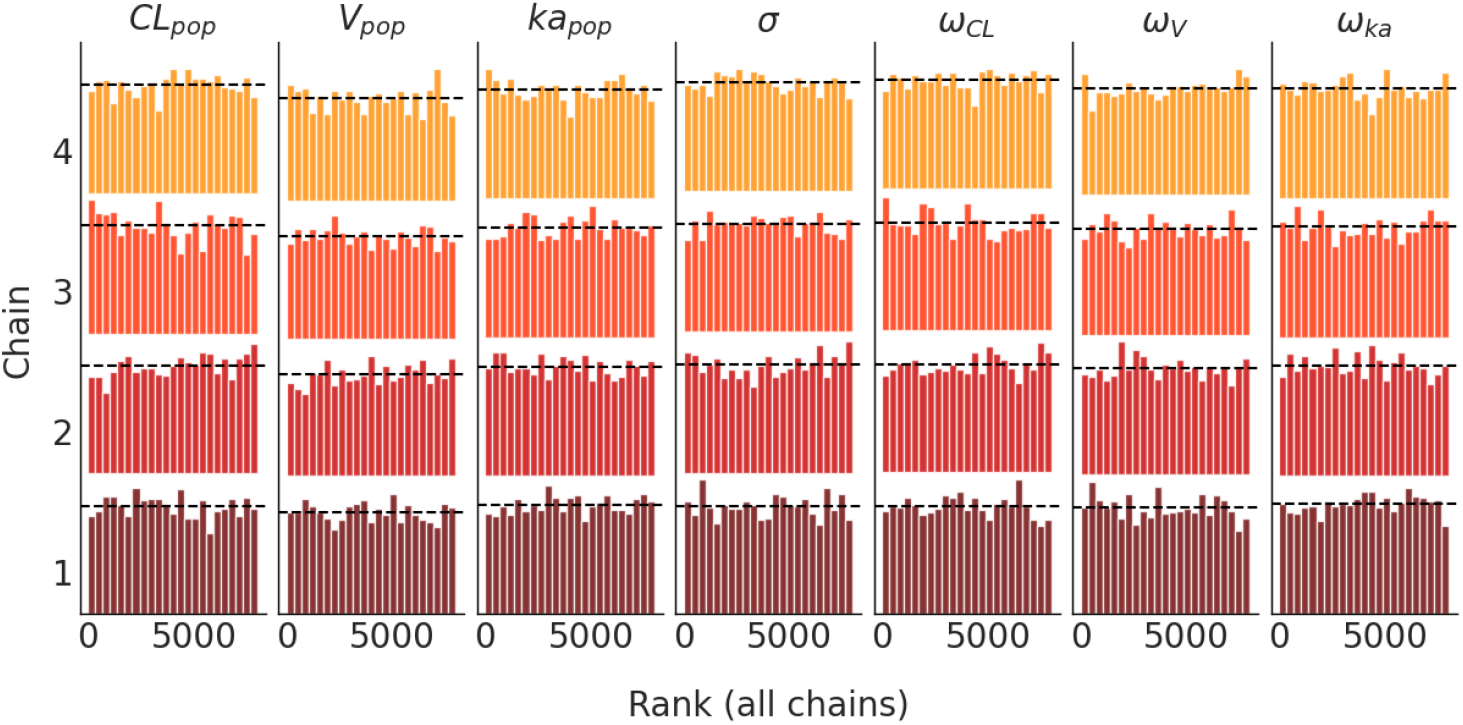
Rank plots (histograms of 2000 ranked posterior draws, ranked over all four chains) of population PK and variability parameters for weakly informative model I fitted against empirical Baclofen data. Rank plots are close to uniform (dashed horizontal lines), thus, the chains mixed well and converged to a stationary distribution.

**Figure C.14.**
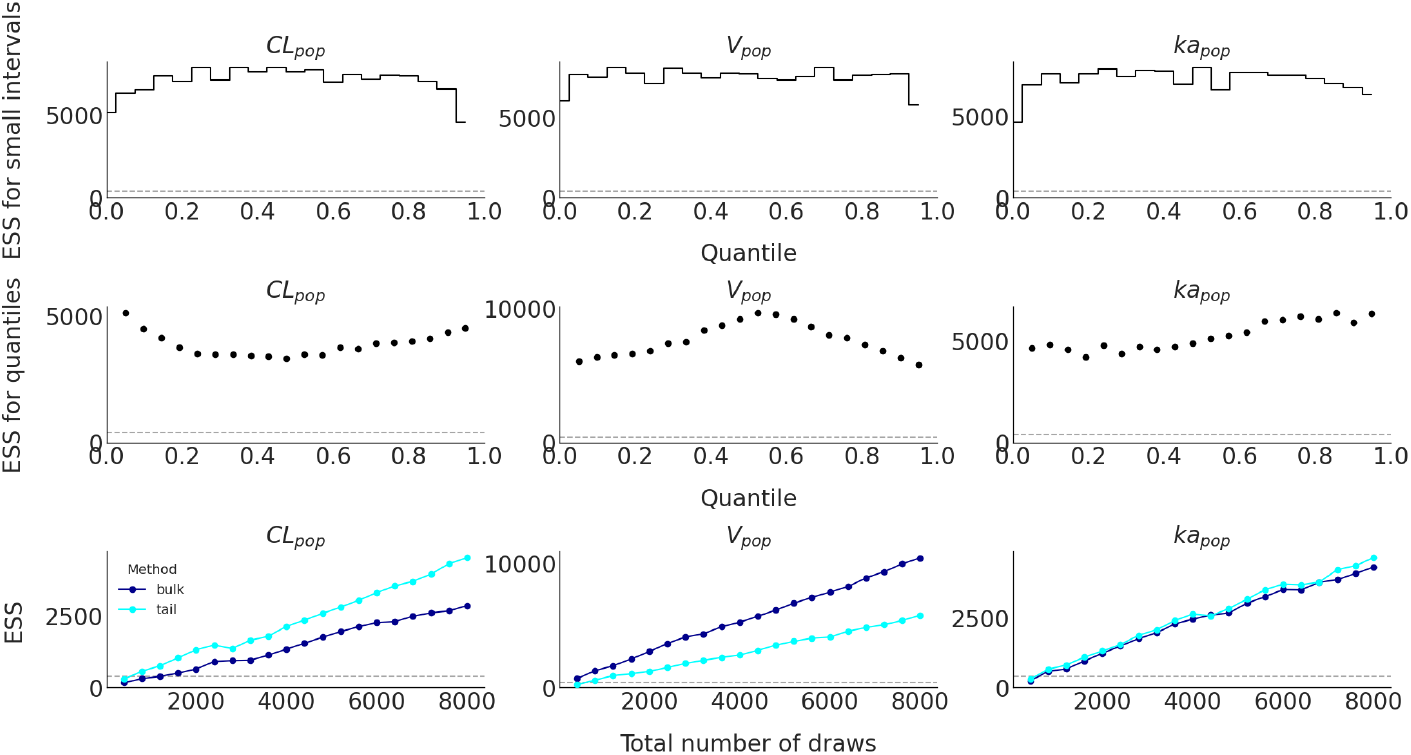
Effective sample size (ESS) plots of population PK parameters using weakly informative model I fitted against empirical Baclofen data. Top: local efficiency of small interval probability estimates, Middle: efficiency of quantile estimates, Bottom: estimated effective sample size (bulk and tail methods) with the increasing number of iterations.

**Figure C.15.**
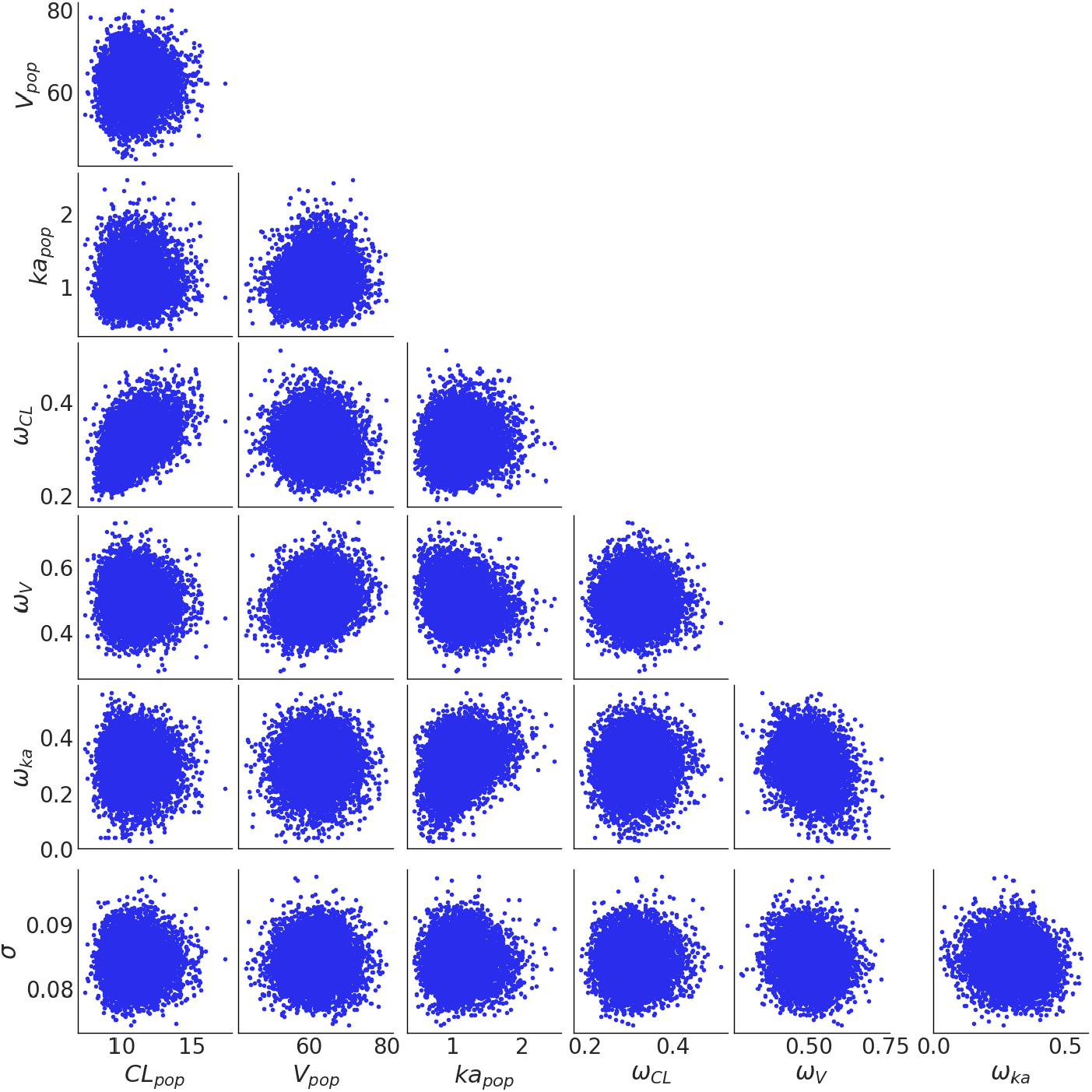
Paired posterior draws (4 chains, 2000 draws each) of population PK parameters using weakly informative model I fitted against empirical Baclofen data.

**Figure C.16.**
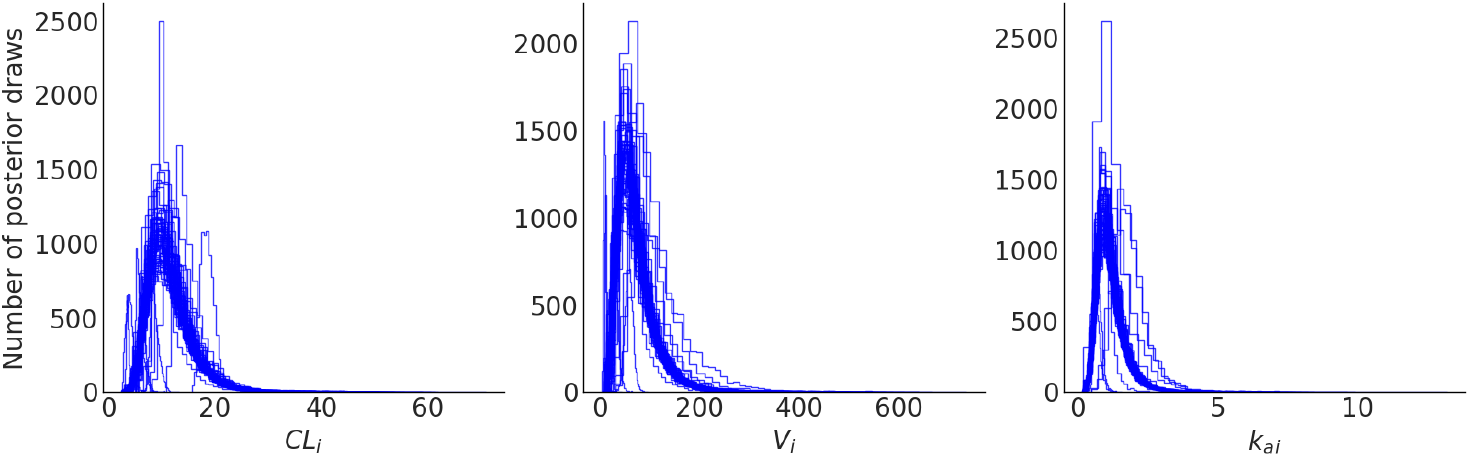
Individual posterior distributions (histograms with 40 bins) of individual one-compartment PK parameters fitted against empirical Baclofen data.

## D. Workflow of the steady state calculation for one-compartment models with Torsten

Translated to algorithm text from the `PredSS_oneCpt.hpp` file of Torsten library^10^.

**Case of interest:** bolus dose in the gut compartment (`cmt = 1`), with `ss = True`.

Set 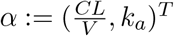

- Computation of amount in gut *y*_*gut*_:

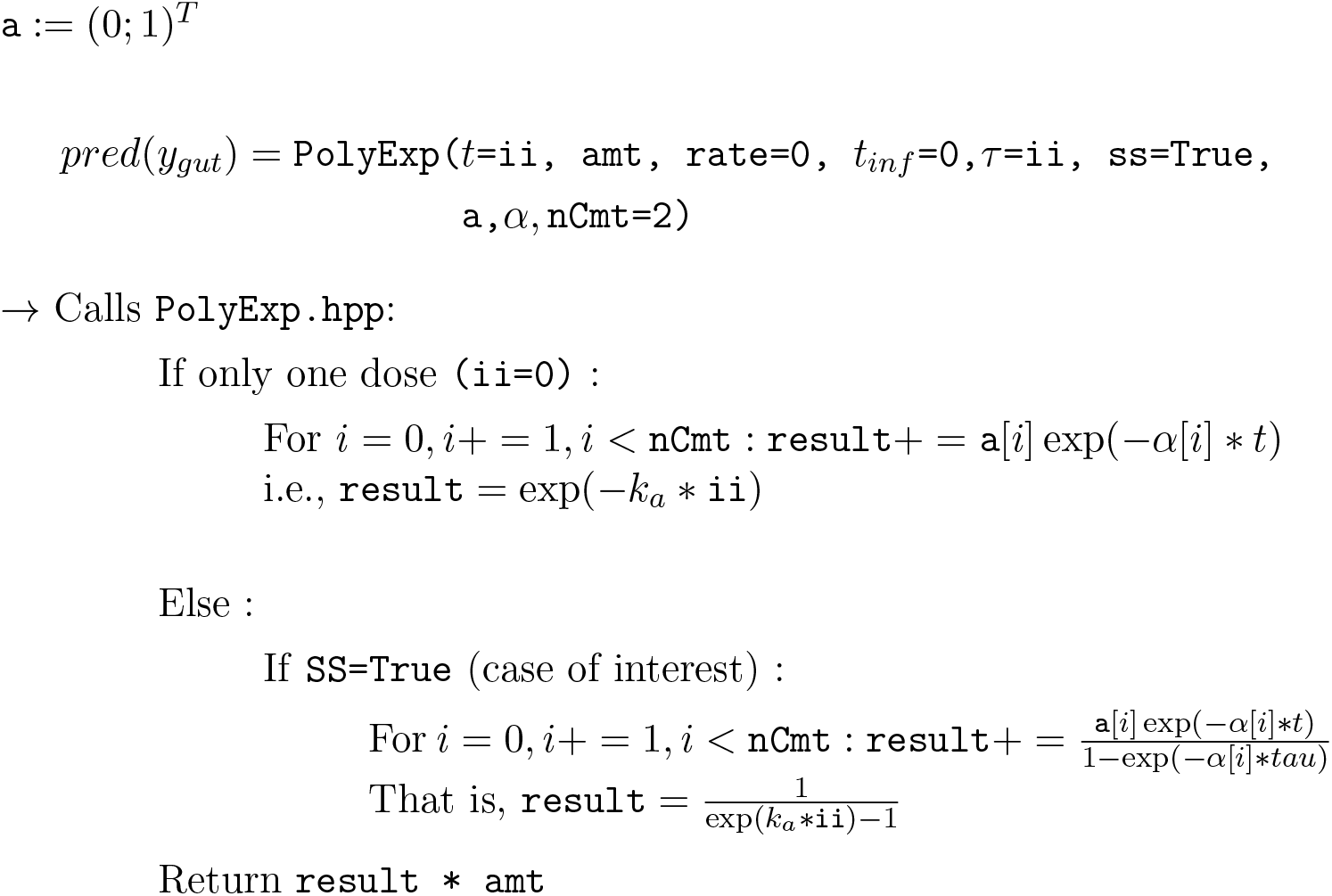
- Computation of amount in central compartment *y*_*cent*_:

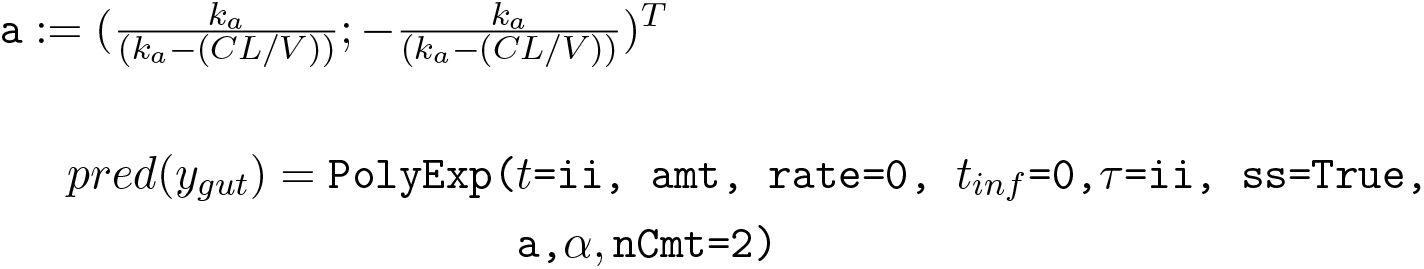

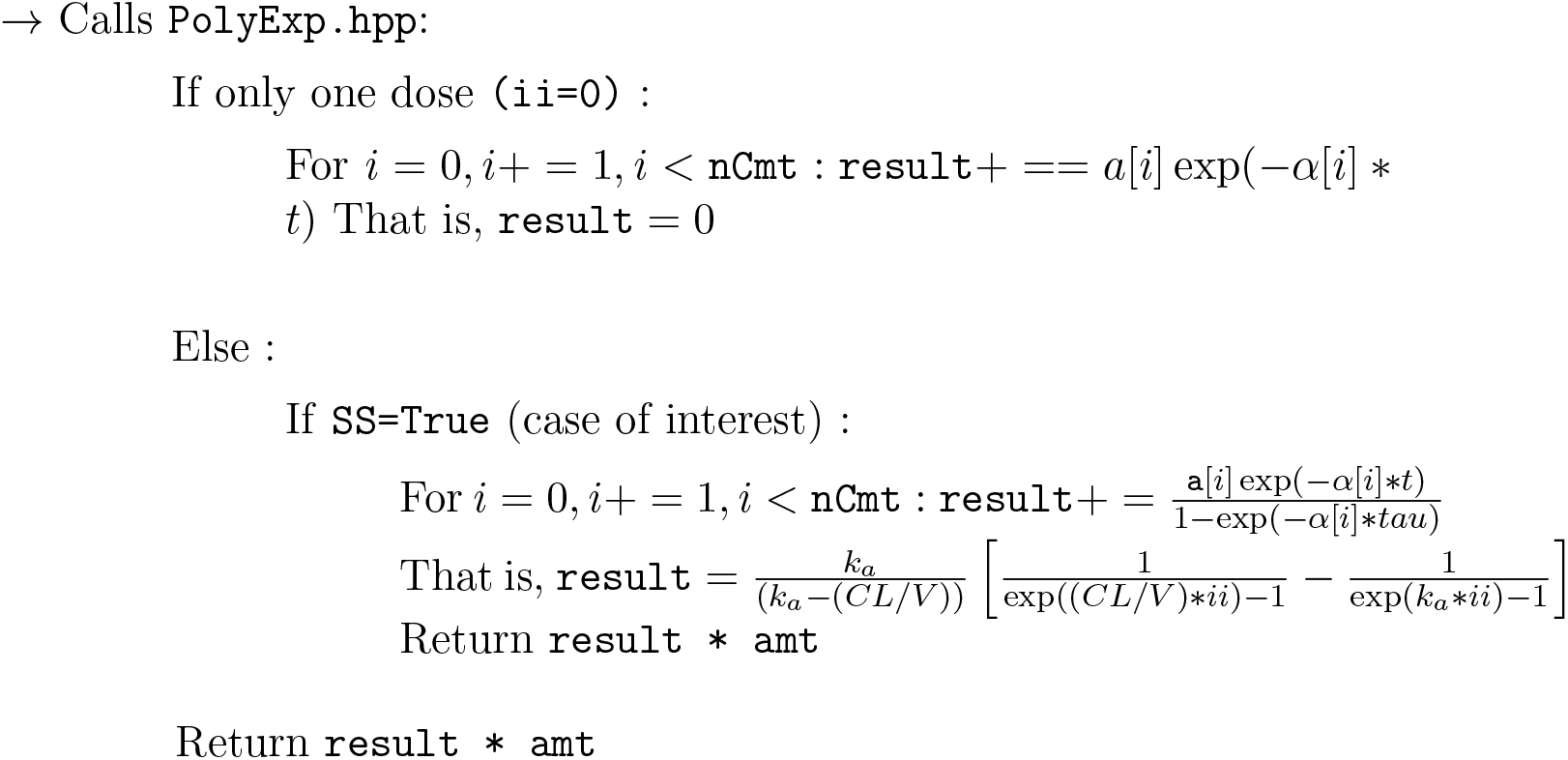

## Notes

### Competing Interest Statement

The authors have declared no competing interest.

